# Understanding the adaptive role of chromosomal inversions across large geographical scales: The potential of pool-seq data

**DOI:** 10.1101/2023.08.20.553987

**Authors:** Anja M Westram, Hernán E. Morales, Kerstin Johannesson, Roger Butlin, Rui Faria

## Abstract

A growing body of research shows that chromosomal inversions, where each arrangement is associated with a certain environment and maintains a set of adaptive alleles, make an important contribution to local adaptation. However, inversions often remain unexplored across large geographical scales. It is unclear whether inversions contribute to adaptation across species ranges, which environmental factors affect arrangement frequencies, and whether the adaptive content of the same arrangement varies between locations. Here, we discuss ideas of how allele frequency data (e.g. pool-sequencing data) can be used to learn about inversions in a simple and cost-effective way. If populations connected by migration differ in arrangement frequency, plotting their SNP allele frequencies against each other will reveal a parallelogram whose corners reflect arrangement frequencies. We demonstrate the usefulness of this approach in locally-adapted populations of the intertidal snail *Littorina saxatilis* (Olivi). For twelve inversions, we estimate arrangement frequencies in 20 populations across the European species range. While roughly half of the inversions contribute to adaptation to the well-studied contrast between wave-exposed and crab-infested habitats, the other half likely contribute to adaptation to shore height. We also find evidence for geographical variation in arrangement content, suggesting variation in adaptive role.

## Introduction

Local adaptation and speciation rely on the evolution of genetic differences between populations. How can such differences arise and be maintained when divergence happens in the face of gene flow, with recombination potentially breaking up favorable allele combinations and homogenizing allele frequencies (Felsenstein 1981)? Theory predicts that genetic architectures that group locally adaptive alleles in the genome or that reduce the negative effects of recombination in other ways will be favored (Smadja and Butlin 2011). Such architectures include chromosomal inversions (Kirkpatrick and Barton 2006; Charlesworth and Barton 2018). Kirkpatrick & Barton’s (2006) model relies on the fact that recombination between the standard and the inverted arrangement in heterokaryotypic individuals (i.e., those with one standard and one inverted arrangement) is largely prevented, so that large “blocks” of alleles are protected from recombination. If the two arrangements contain multiple alleles adapted to alternative environments, inversions are therefore favored under divergent adaptation with gene flow.

Recent empirical genomics work supports these theoretical predictions. For example, cod populations in fjords are likely to be adapted to that particular environment via a chromosomal rearrangement containing alleles beneficial in low salinities (Barth et al. 2017). Adaptive differences between annual and perennial ecotypes of *Mimulus* monkeyflowers can be traced to a chromosomal inversion (Twyford and Friedman 2015), and inversions are associated with marine-freshwater divergence in stickleback fish (Jones et al. 2012).

While the role of inversions in local adaptation is becoming clearer, it is less well understood how this plays out on large geographical scales. Many taxa have wide geographical distributions, where similar environmental contrasts can occur repeatedly, but where selective factors can also be combined differently in different geographical locations. Then, the question is whether the same arrangements are repeatedly involved in adaptation and whether they are consistently associated with specific environmental factors. Inversions containing multiple co-adapted alleles might represent efficient “transport vehicles” between locations with similar environments (Westram et al. 2022). On the other hand, because of recombination restriction, arrangement allelic content is somewhat inflexible, and difficulties in generating the “right” allelic combinations might prevent their repeated use when multiple environmental factors vary in complex environments (Roesti et al. 2022).

Some studies have found evidence for the re-use of inversions in adaptation across multiple locations: In *Drosophila subobscura*, the same inversions show repeated large-scale latitudinal clines on different continents and are associated with temperature adaptation (Kapun et al. 2014); in honeybees, the same inversions are repeatedly associated with altitude adaptation in different mountain regions (Christmas et al. 2019); and in seaweed flies, inversions show similar environmental associations in Europe and North America (Mérot et al. 2018). However, studies dissecting the adaptive role of inversions on large geographical scales are still relatively rare, especially in non-model species.

One advantage of gathering data on inversions across large geographical scales is that this might allow to disentangle different potential selective factors. Locally, multiple potential selective factors are often associated in space, e.g., different climate variables and biotic factors such as interacting species. In geographically distant sites, environmental factors might be associated in different ways, making it possible to identify those that are statistically associated with arrangement frequencies.

A directly related question is how arrangement content varies across space. Large inversions can contain large numbers of functional loci, and, within the same arrangement (e.g., the inverted arrangement), the genotypes at these loci may differ between geographical locations. The same arrangement could therefore also contribute to adaptation in different ways. For example, take the hypothetical example of a plant species adapted to different altitudes. If the same inversion contributes to altitude adaptation in two locations, the causal SNPs are not necessarily the same. The same inversion, if containing different adaptive alleles, could even contribute to altitude adaptation in one location and to herbivore adaptation in another. In other words, inversions are a useful evolutionary “toolbox” for local adaptation which, could be “filled” with different contents of adaptive alleles (i.e., different tools), depending on locally available genetic variation and local selection pressures.

Studying the adaptive role of inversions on large geographical scales therefore requires information about arrangement frequencies as well as information about SNP allele frequencies within arrangements for multiple locations. Recent studies have used individual next-generation sequencing data for inversion detection and genotyping via elevated linkage disequilibrium, caused by the reduction of recombination between arrangements (Kemppainen et al. 2015; Faria et al. 2019a; Huang et al. 2020; Mérot 2020). Individual sequencing data also provides information on inversion content. However, when including many locations this approach incurs high costs. In other population genetics contexts, pool-sequencing data, where multiple individuals from the same population are sequenced together without individual barcoding (Schlötterer et al. 2014), represents a cost-effective alternative to individual sequencing. This type of data provides population-level SNP allele frequency estimates but no individual genotypes and, therefore, cannot be used in the LD-based approaches mentioned above. F_ST_ scans with pool-seq data can provide indirect evidence for inversions under divergent selection via the presence of large blocks of elevated F_ST_ between populations (e.g. Gould et al. 2017; Morales et al. 2019). However, to our knowledge, pool-seq data alone could so far not be used to determine arrangement frequencies, because the association between F_ST_ and arrangement frequencies is not straightforward. Determining arrangement frequencies in pool-seq data has therefore required prior knowledge about arrangement-diagnostic SNPs (i.e. SNPs that are fixed different between the two arrangements) (Kapun et al. 2014).

In this paper, we illustrate the use of pool-seq data for studying inversions across large geographical scales. We show that some LD information is retained in pool-seq data (or SNP allele frequency data from other sources), beyond the short-range LD within sequencing reads, and can be used to detect inversions *de-novo*, determine population arrangement frequencies, and study within-arrangement polymorphisms (i.e., SNPs that vary between copies of the same arrangement). In the first part of the paper, we explain our approach conceptually. It relies on standard plots of allele frequency data; we aim at emphasizing the simplicity of applying this approach and therefore do not provide a specific data analysis pipeline or software. Our approach requires, at a minimum, SNP allele frequency data from two diverging populations connected by gene flow and becomes especially useful when extended to multiple population pairs from the same taxon. We then use these ideas to test the role of inversions in local adaptation in the rough periwinkle *Littorina saxatilis* (Olivi), a marine snail, across large geographical scales, demonstrating the role of inversions in parallel evolution and showing within-arrangement differences between locations.

## Rationale

### Determining arrangement frequencies from SNP allele frequency data

When two populations diverge with gene flow, a plot of SNP allele frequencies in population 1 against frequencies in population 2 typically shows a correlation, with points scattered around the identity line (where x=y) (Fig. 1A). When the two populations differ in arrangement frequency for an inversion, allele frequencies will still be correlated (because within each arrangement, gene flow and recombination between populations take place), but the correlation lines will be shifted from the identity line due to the different arrangement frequencies. This generates a “parallelogram” pattern that can be used to detect inversions and determine arrangement frequencies. We explain this pattern in the following.

**Fig. 1:**
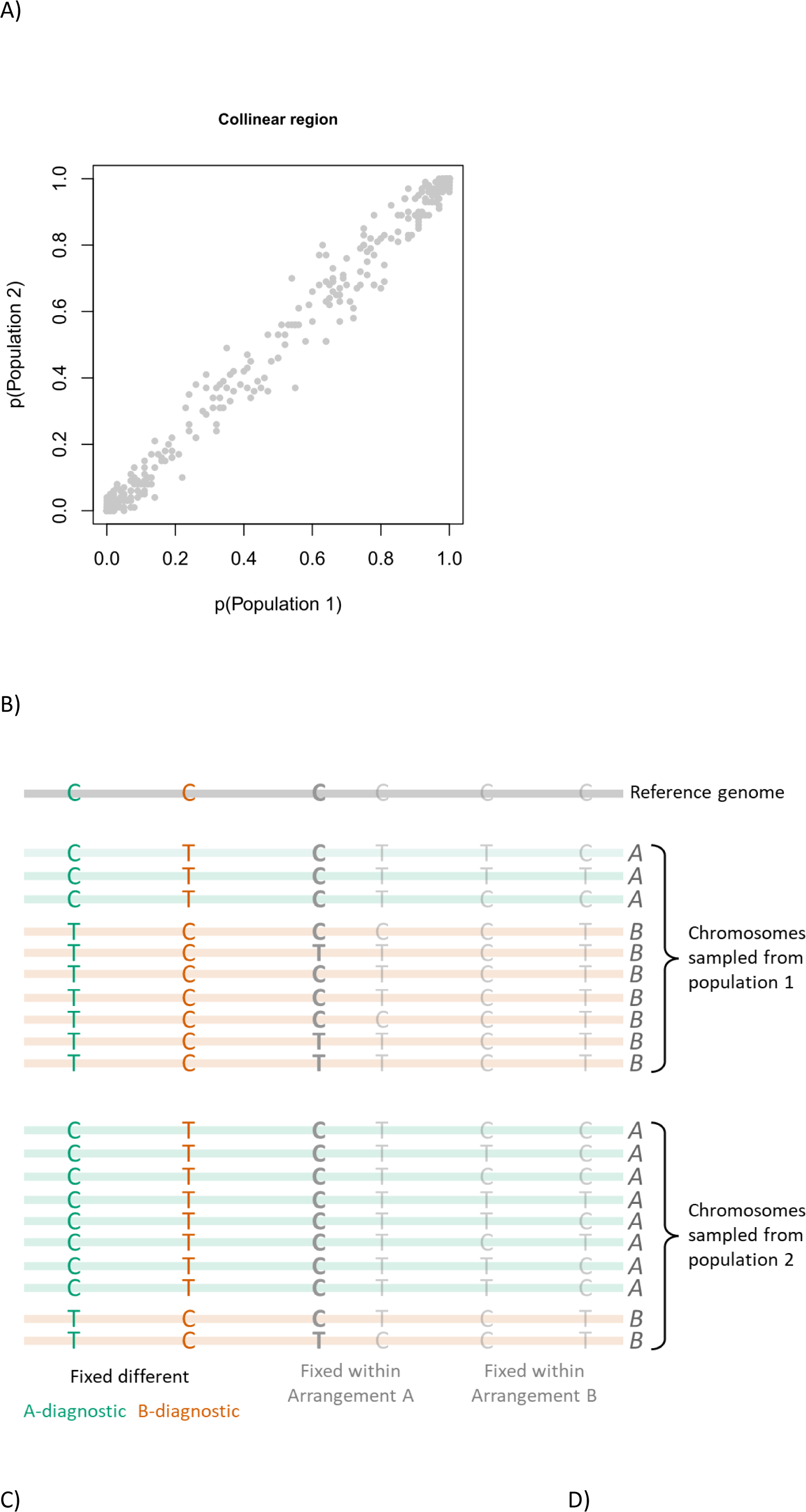
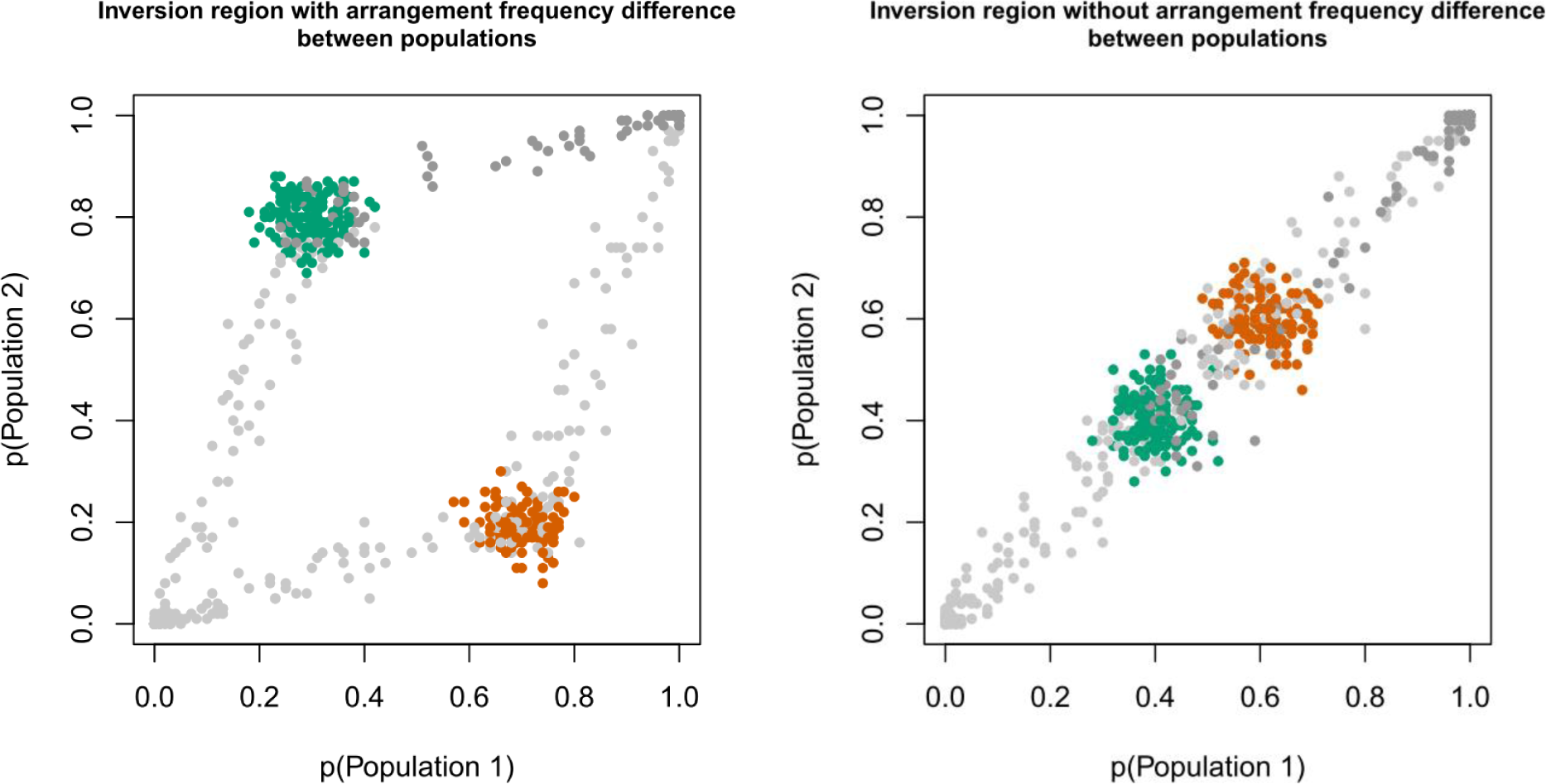
The emergence of parallelograms in allele frequency plots. A) Simulation of a collinear region. B) Illustration of a chromosomal region with inversion polymorphism (light green horizontal line: arrangement A; light orange line: arrangement B; shown in the same orientation for illustrative purposes). The reference genome (top) is assumed to have a C allele at each SNP position for simplicity. SNPs fixed different between arrangements might have the reference allele either in arrangement A (A-diagnostic SNPs, green) or arrangement B (B-diagnostic SNPs, orange). Alternatively, SNPs can be fixed in only one arrangement (grey). C) Simulation with *p*_*A*,1_ = 0.3 and *p*_*A*,2_ = 0.8 (as in panel B). D) Simulation with *p*_*A*,1_ = *p*_*A*,2_ = 0.4. In C) and D), SNPs considered arrangement-diagnostic are indicated in green (A-diagnostic) and orange (B-diagnostic). The SNP type variable in one arrangement discussed in the main text is shown in bold in B) and by darker grey points in C) and D). Details about simulations in Supplementary Text.

We assume two populations connected by levels of gene flow high enough to generate correlations in allele frequencies in the background genome (e.g., Fig. 1A). We focus on a genomic region with a simple inversion polymorphism; i.e., two alternative arrangements, labelled A and B. It is irrelevant here which arrangement is the ancestral and which is the inverted arrangement. Arrangements A and B segregate at potentially different frequencies in the two populations. Thus, there are four arrangement frequencies, *p*_*A*,1_, *p*_*A*,2_, *p*_*B*,1_ = 1 − *p*_*A*,1_, and *p*_*B*,2_ = 1 − *p*_*A*,2_.

Unless the inversion is very young, some SNPs will be (nearly) differentially fixed between the two arrangements (Fig. 1B, green and orange). These SNPs provide direct estimates of arrangement frequencies (Fig. 1C). If we assume allele frequencies are given in relation to a reference genome that sequencing data were mapped to, there are two sets of such diagnostic SNPs: SNPs where the A arrangement is fixed for the reference allele (“A-diagnostic SNPs”, green), and SNPs where the B arrangement is fixed for the reference allele (“B-diagnostic SNPs”, orange). The frequencies of A-diagnostic SNPs in population 1 and population 2 provide estimates of the A arrangement frequencies in these two populations (*p*_*A*,1_ and *p*_*A*,2_), and thus also of *p*_*B*,1_ = 1 − *p*_*A*,1_and *p*_*B*,2_ = 1 − *p*_*A*,2_. Frequencies of B-diagnostic SNPs can be used in a similar way. These arrangement-diagnostic SNPs are also the strongest candidate loci for local adaptation – especially if they are diagnostic across multiple locations – as they achieve the highest differentiation between environments when the inversion is divergently selected.

SNPs polymorphic in both arrangements require recent exchange between arrangements, or repeated mutation, and are assumed to be rare. However, some SNPs will be fixed in arrangement A but polymorphic in arrangement B, or vice versa (Fig. 1B). These SNPs are not directly indicative of arrangement frequencies. However, they can only obtain a restricted range of allele frequencies. To give an example, we here focus on SNPs where arrangement A is fixed for the reference allele. We assume arrangement A has a frequency of *p*_*A*,1_ = 0.3 in population 1 (Fig. 1B). In this case, all these SNPs must have a minimum allele frequency of 0.3 in population 1; each SNPs has an allele frequency between 0.3 and 1, depending on its frequency within arrangement B (one example shown in Fig. 1B, bold grey SNP). The same argument applies to population 2. Because there is gene flow between populations within arrangements, allele frequencies will be correlated between populations; but the correlation line will start at the minimum allele frequencies (*p*_*A*,1_, *p*_*A*,2_) (Fig. 1C, dark grey circles).

Mathematically, the situation for a SNP fixed for the reference allele in arrangement A, but polymorphic in arrangement B, is as follows. The expected frequency for SNP *i* is

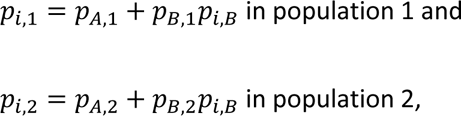

where *p*_*i*,*B*_ is the frequency of the reference allele for this SNP within arrangement B. If gene flow is high, *p*_*i*,*B*_ is expected to be roughly the same in both populations, and thus the two equations can be combined to express the correlation between the allele frequencies in the two populations:

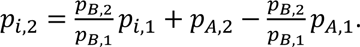

This equation confirms that the line relating the allele frequencies in population 1 and 2 goes through (*p*_*A*,1_, *p*_*A*,2_) and through (1,1), as explained in the example above, and shows that the slope is 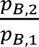 (Fig. 1C, dark grey circles).

Analogous lines will form for the remaining three sets of SNPs (fixed for the alternative allele in A, variable in B; fixed for the reference allele in B, variable in A; fixed for the alternative allele in B, variable in A) (Fig. 1B,C). Together, these four lines form a parallelogram, whose corners are at (0,0), (1,1), (*p*_*A*,1_, *p*_*A*,2_) and (*p*_*B*,1_, *p*_*B*,2_) (Fig. 1C).

If arrangement frequencies do not differ between populations, no parallelogram will form as it “collapses” into a line (Fig. 1D). However, if arrangement-diagnostic SNPs are known, arrangement frequencies can still be determined (colored points in Fig. 1D).

The equations shown here assume no noise in the data and that within-arrangement frequencies are the same in population 1 and 2, leading to a perfect parallelogram. In empirical data, statistical sampling (included in Fig. 1) generates noise when sampling individuals from the population and pooling potentially slightly unequal amounts of DNA. In addition, due to evolutionary sampling (i.e. genetic drift), allele frequencies within arrangements will differ between populations, partly depending on the level of gene flow. Importantly, both sources of noise will generate scatter around the parallelogram but should not generate a bias or distortion of the shape.

Some rare SNPs may violate our assumptions: SNPs could be variable in both arrangement A and arrangement B, and therefore be located inside the parallelogram. SNPs could also show allele frequency differences within arrangements between populations (potentially due to selection) and could thus in principle be located anywhere in the plot. However, these issues are expected to affect a minority of the SNPs in an inversion region, and they will again not distort the overall parallelogram shape.

Thus, we expect that in many systems it should be possible to detect inversions by the presence of parallelograms in allele frequency plots, and to estimate arrangement frequencies by reading off the allele frequencies at the parallelogram corners.

## Study system

To use the above ideas for studying inversion evolution on large geographical scales, we focused on a system with known inversion polymorphism, the marine snail *Littorina saxatilis,* that shows parallel divergent adaptation in many geographical locations. *L. saxatilis* occurs on rocky shores across Europe and North America. The two most well-studied ecotypes of *L. saxatilis* are the “Crab” ecotype - large, thick-shelled, wary snails adapted to crab predation - and the “Wave” ecotype - small, thin-shelled, bold snails adapted to wave exposure. Both ecotypes occur across Europe and have probably evolved repeatedly at least in Spain, Sweden, and the UK (Butlin et al. 2014). The ecotypes are locally connected by gene flow and form hybrid zones where their habitats adjoin (Westram et al. 2021).

In addition, *L. saxatilis* also shows local adaptation to different shore levels (Johannesson et al. 1995). High shore levels are associated with increased air and sun exposure compared to low shores, and thus high-shore snails experience more extreme temperature and higher desiccation risk. Importantly, Crab-Wave and Low-High adaptation are not independent (explained in detail in Supplementary Text; Morales et al. 2019), as high shore levels correspond to Crab habitat and low shore levels correspond to Wave habitat in Spain, while this pattern is reversed in the UK and France (Fig. 2). In Sweden, both Crab and Wave occur across both shore levels (Fig. 2).

**Fig. 2:**
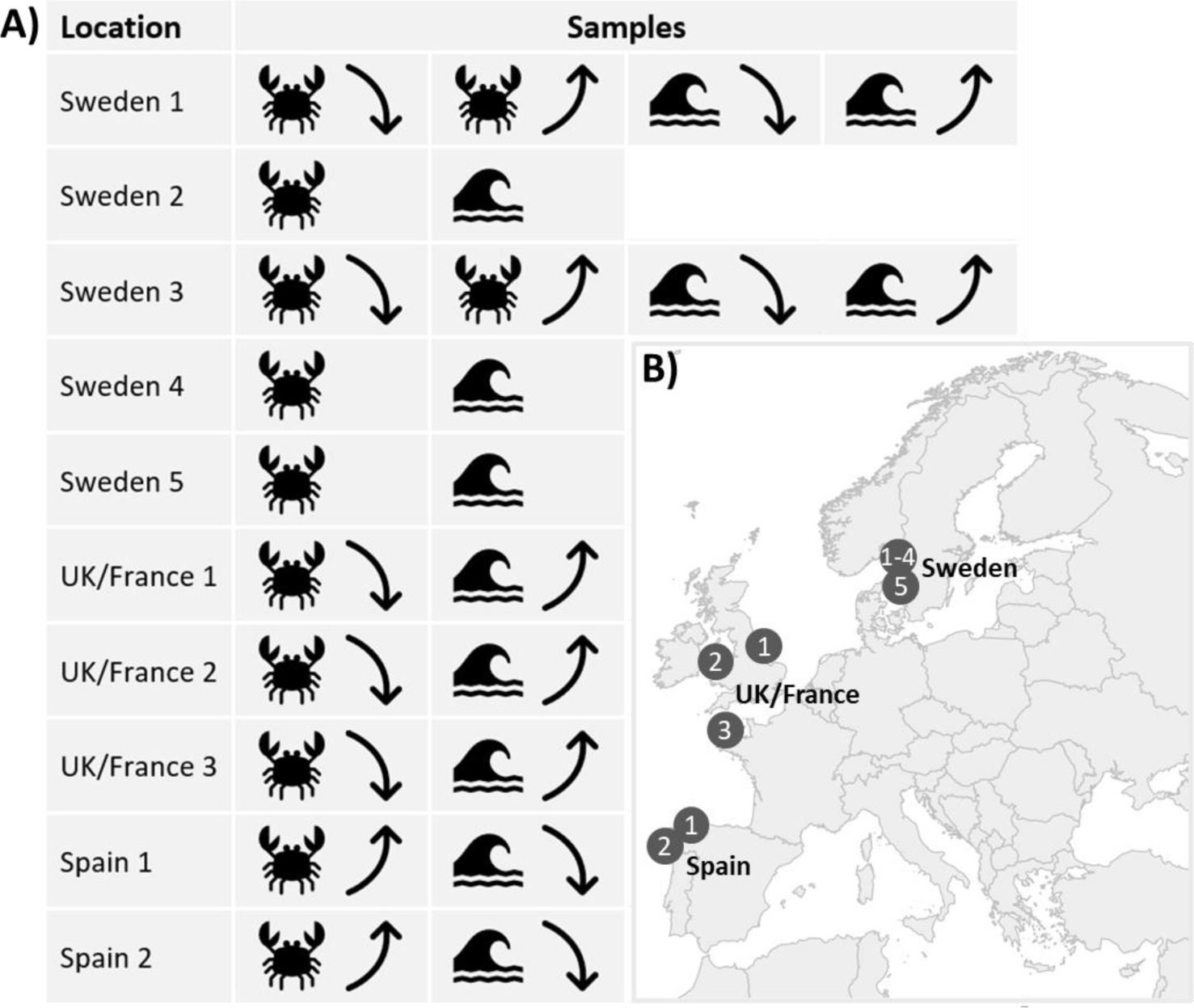
Overview of the samples (pools) used in this study (see also Morales et al. 2019). A) Up to four samples were obtained in each of ten geographical locations. Crab and Wave ecotype are indicated by symbols; Low and High are indicated by an upward and a downward arrow, respectively. Note that the association between the Crab-Wave and the Low-High axis varies between geographical areas. For Sweden 2, 4 and 5, pools contained individuals sampled randomly across both shore levels. B) Partial map of Europe showing the ten sampling sites.

Previous work on the Swedish west coast has shown that the Crab and Wave ecotype differ in arrangement frequency for multiple chromosomal inversions (Faria et al. 2019, Westram et al. 2021) and that divergent traits map to these inversions (Koch et al. 2021). A study based on pool-seq from Spain, France, the UK, and Sweden found that multiple genomic regions corresponding to polymorphic inversions in Sweden show high levels of differentiation (e.g., elevated F_ST_) between Crab and Wave ecotypes in other European locations as well (Morales et al. 2019). Some of the inversion regions also showed elevated differentiation between the low and high shore (Morales et al. 2019).

While these results suggest a role of inversions in parallel evolution in *L. saxatilis*, three important aspects are not well understood. 1) The arrangement frequencies are only known for four Swedish locations and could thus far not be determined for other European locations due to the use of pool-seq data. In particular, for locations where F_ST_ is low between Crab and Wave or Low and High pool-sequencing samples, it is unclear whether this is due to the absence of inversion polymorphism or due to similar arrangement frequencies in both populations. 2) As a consequence, the association of different arrangements with the different selective axes could be addressed only indirectly (Morales et al. 2019) and needs confirmation with actual arrangement frequency data. 3) It is unclear whether the adaptive alleles inside arrangements differ between geographical locations.

In this paper, we address these three points, using pool-seq data according to the ideas explained above, with three main aims: 1) Determining arrangement frequencies in multiple geographical locations across Europe; 2) Testing whether inversions are consistently associated with Crab-Wave or Low-High selection; 3) Analyzing the allelic content of different arrangements and comparing it between locations. We confirm our inferences of arrangement frequencies with individual sequencing data.

## Methods

### Sampling and pool-sequencing

We reused the whole-genome pool-seq data published by Morales et al. (2019). Briefly, snails were collected from 10 locations in three geographical areas - Sweden, UK/France, and Spain (Fig. 2, Table S1). UK and France are grouped together here as they are geographically relatively close and share the same shore distribution of ecotypes (Fig. 2). Snails were collected from Crab and Wave habitat as well as from the Low and High shore (i.e., the lowest and highest part of the local distribution).

In Spain and UK/France, the Crab-Wave and the Low-High axes are associated, but with different directionalities (Fig. 2). In these cases, there are only two possible types of samples (Crab-Low and Wave-High in UK/France; Crab-High and Wave-Low in Spain), each of which is represented by a pool of 100 females. In Sweden, both the Crab and the Wave ecotype can occur on the low and high shore (in separate parts of the shore), so that four different types of samples are possible. For two locations (Sweden 1 and 3), we generated pools representing all four possible combinations, each containing 50 females. When doing Crab-Wave comparisons for these locations (see below), we combined allele counts for the Low and High sample for each of Crab and Wave. For the remaining three Swedish locations (Sweden 2, 4 and 5), only Crab and Wave ecotypes were collected randomly across shore levels. Each of these pools contained DNA from 24 individuals (including both sexes).

For each pool (“sample” in the following), tissue of the different individuals was combined (i.e., pooled), DNA was extracted, genomic libraries were generated and sequenced on an Illumina HiSeq2500 machine. Reads for each sample were trimmed and mapped to the *L. saxatilis* reference genome. After further bioinformatic processing, we obtained allele count data for each sample and used these to calculate allele frequencies. For details of the pooling procedure and bioinformatic processing, see Morales et al. (2019). We only retained biallelic SNPs with a coverage depth of at least 25x in each pool and a minor allele frequency of 0.02 across all pools together.

### De-novo detection of inversion polymorphism

In this study, we focus mainly on inversion regions described previously by Westram et al. (2018) and Faria et al. (2019) for the Swedish west coast, determining arrangement frequencies in other European locations (next section). However, as we mention above that pool-seq data might be used for inversion de-novo detection as well, we also ask whether de-novo detection would have been possible with our data.

We generated allele frequency plots analogous to Fig. 1C / Fig. 1D for each Crab-Wave pair in 4cM windows along the genome. 4cM were used to ensure enough SNPs in most windows. One of us, unaware of chromosome and location IDs, evaluated each plot, noting in which window a parallelogram was visible (“Yes”, “No”, or “Uncertain”). For this, each set of windows belonging to the same chromosome was kept intact, as seeing differentiation in the collinear background is necessary to evaluate the presence of a parallelogram; but the chromosome and location IDs were removed, and the order of the sets was randomized. We then compared the results of the scoring with the locations of known inversions.

### Estimation of arrangement frequencies

Our main goal was to estimate arrangement frequencies. We included 12 inversion regions on 9 different chromosomes from Faria et al. (2019), excluding arrangements on linkage group 14, which showed inconsistent patterns between Spain and the other countries and might contain complex or overlapping rearrangements, and regions on linkage group 12, which is probably a sex chromosome (Hearn et al. 2022). The inversions are labelled LGC1.1, LGC1.2, LGC2.1, LGC4.1, LGC6.1/2, LGC7.1, LGC7.2, LGC9.1, LGC10.1, LGC10.2, LGC11.1, and LGC17.1, where the first number in each label refers to the linkage group the inversion region is located in. We used the chromosomal coordinates from Westram et al. (2021; innermost coordinates).

As shown in Fig. 1C, arrangement frequencies could be estimated for a single Crab-Wave or Low-High sample pair from a single location. Here, as we sampled multiple locations, we first identified “globally arrangement-diagnostic SNPs” (SNPs that were arrangement-diagnostic across many locations) and used only those to estimate arrangement frequencies. This approach has three advantages: 1. It ensures that the “A arrangement” and the “B arrangement” are labelled consistently across locations, as the same diagnostic SNPs are used; 2. It improves the detection of the parallelogram corners for locations where the allele frequency data are noisy; 3. It allows for the estimation of arrangement frequencies in locations where the frequency does not differ between populations (i.e. where the parallelogram collapses into a line) (Fig. 1D).

To determine A- and B-diagnostic SNPs for each known inversion region, we performed three steps, explained in detail in the Supplementary Text. Briefly, we first plotted allele frequency data for all SNPs from the inversion region analogous to Fig. 1C / Fig. 1D for each Crab-Wave population pair and ensured visually that multiple pairs showed a parallelogram pattern (this was the case for all studied inversions). We then identified the SNPs with the highest F_ST_ (>80% quantile of the F_ST_ distribution) for each pair. These SNPs roughly correspond to the arrangement-diagnostic SNPs (i.e., the SNPs in the (*p*_*A*,1_, *p*_*A*,2_) and (*p*_*B*,1_, *p*_*B*,2_) corners of the parallelogram). The reason is that differentiation between populations with different arrangement frequencies is highest for those SNPs that are fixed different between arrangements (again assuming no large allele frequency differences within arrangements between populations, which is appropriate for the majority of SNPs). As we were looking for globally arrangement-diagnostic SNPs, we then retained only those SNPs as candidates that were in the high-F_ST_ class in at least 50% of the included locations. Finally, we ensured to retain only SNPs whose allele frequencies are strongly associated with arrangement frequencies across locations. For that, we ran a PCA for all populations, including all candidate SNPs. The first axis of this PCA sorts populations roughly by arrangement frequency. The SNPs that have the greatest impact on this axis are the ones most strongly associated with arrangements across locations. We therefore retained only those candidate SNPs above the 90% quantile of the distribution of absolute loadings on PC1.

Each inversion arrangement might be associated with the reference genome allele at some SNPs and with the alternative allele at others (Fig. 1). For that reason, some SNPs have positive loadings on PC1 while others have negative loadings. We therefore obtained two sets of arrangement-diagnostic SNPs, “A-diagnostic SNPs” (positive loadings; reference allele associated with the arrangement arbitrarily labelled “A”) and “B-diagnostic SNPs” (negative loadings; reference allele associated with arrangement B).

Having obtained two sets of globally diagnostic SNPs for each inversion, we calculated an estimate of the frequency of arrangements A and B for each sample shown in Fig. 2 by averaging across the frequencies of the diagnostic SNPs in that sample.

The inversion on linkage group 6 (labelled LGC6.1/2) contains a part where most likely a second inversion has occurred, overlapping the original, larger inversion. Therefore, three mostly non-recombining haplotypes exist (Faria et al. 2019a). In that case we did not expect to observe a simple parallelogram when including SNPs from the whole inversion region. We therefore restricted all analyses of this inversion to the region affected by only a single inversion, which contains only two haplotypes, applying a 1cM buffer (0-7.73cM).

We validated our arrangement frequency results using individual sequencing data. A recent study obtained capture sequencing data for hundreds of individuals using probes randomly distributed throughout the genome for two of the Swedish locations included here (Sweden 3 and Sweden 4). As multiple probes covered each inversion, reliable arrangement frequency estimates could be obtained using an LD-based method (Westram et al. 2021, Faria et al. 2019). Here, we compared the arrangement frequency estimates obtained in the two studies.

### Association of arrangements with selective axes

We first tested, for each inversion and geographical location, whether an inversion showed elevated differentiation between two samples (i.e., the Crab and Wave sample or the Low and High sample). We used the empirical distribution of SNP F_ST_ values as the null distribution, considering only genomic regions outside inversions. We considered an inversion region as highly differentiated if the F_ST_ calculated from the estimated arrangement frequencies was above the 95% quantile of this distribution. However, because arrangement frequencies and SNP frequencies are not directly comparable and were obtained using different methods, the results should be interpreted as a general indication of high differentiation rather than a thorough test for divergence under selection.

Frequency differences between the two samples from the same location in Spain or UK/France could reflect a response to either the Crab-Wave or the Low-High selective axis. The distinction can only be made when including samples from multiple geographical areas where the two selective axes are associated in different ways (Fig. 2). For each inversion, we therefore ran a Wilcoxon signed rank test including all Crab-Wave pairs to test whether one arrangement is significantly more common in one ecotype. We performed the analogous test for the Low-High pairs.

### Analysis of arrangement content

Our aim here was to test whether the same arrangement might contain different adaptive alleles in different geographical areas. If that was the case, one would expect that SNPs that are arrangement-diagnostic in some locations may show different patterns in other locations. First, within each geographical area, we identified SNPs that show a (nearly) fixed difference between the two arrangements (“locally arrangement-diagnostic SNPs”). For that, for each SNP, we calculated the Euclidean distance between a vector of the allele frequencies for the focal SNP and a vector of the frequencies of arrangement A, including all populations in the focal geographical area. We did the same for a vector of the frequencies of arrangement B and retained the smaller Euclidean distance (as the SNP could be either A- or B-diagnostic). We then selected the SNPs with the Euclidean distance below the 0.01 quantile of the distribution of Euclidean distances. We visualized the associations of these SNPs with the inversion in all geographical areas by plotting the regression line (lm() function in R) of SNP frequency against arrangement frequency for each area separately. If a SNP contributes to adaptation only locally, we expect that it will not be strongly associated with the inversion in other geographical areas. If a SNP contributes to adaptation in multiple locations, but does not contribute to adaptation to the same selective axis as the inversion, its association with the inversion might be reversed between locations. For example, if arrangement A contributes to Crab adaptation and arrangement B to Wave adaptation, alleles contributing to high shore levels in this inversion would be found in arrangement A in Spain, but in arrangement B in the UK. We note that, of course, the existence of these expected patterns does not necessarily indicate that the respective SNPs directly contribute to adaptation – they might be neutral SNPs in LD with adaptive SNPs, or they might show the pattern purely by chance. Our analysis is simply aimed at testing whether location-specific arrangement contents exist in general.

The regression between SNP and inversion frequency is most informative when inversion frequency varies strongly between populations. We therefore performed this analysis only on two arrangements showing clear associations with one environmental axis each (LGC6.1/2: Crab-Wave axis; LGC9.1: Low-High axis).

All analyses were performed in R version 4.2.1 (R Core Team 2021).

## Results

### Allele frequencies and de-novo detection of inversion polymorphism

Allele frequencies were generally correlated between samples within locations (Fig. S1), but some genomic regions showed visually identifiable parallelograms. We visually detected parallelograms in at least two locations for each of the inversion regions known from Faria et al. (2019) (Fig. S1, Fig. S2). Allele frequencies for one inversion region are shown as an example in Fig. 3. The corners and edges of the parallelograms were often clearly visible by eye; however, some parallelograms were quite noisy, and there were always some SNPs “inside” the parallelograms, not clearly associated with the parallelogram corners or edges (Fig. 3, Fig. S1, Fig. S3).

**Fig. 3:**
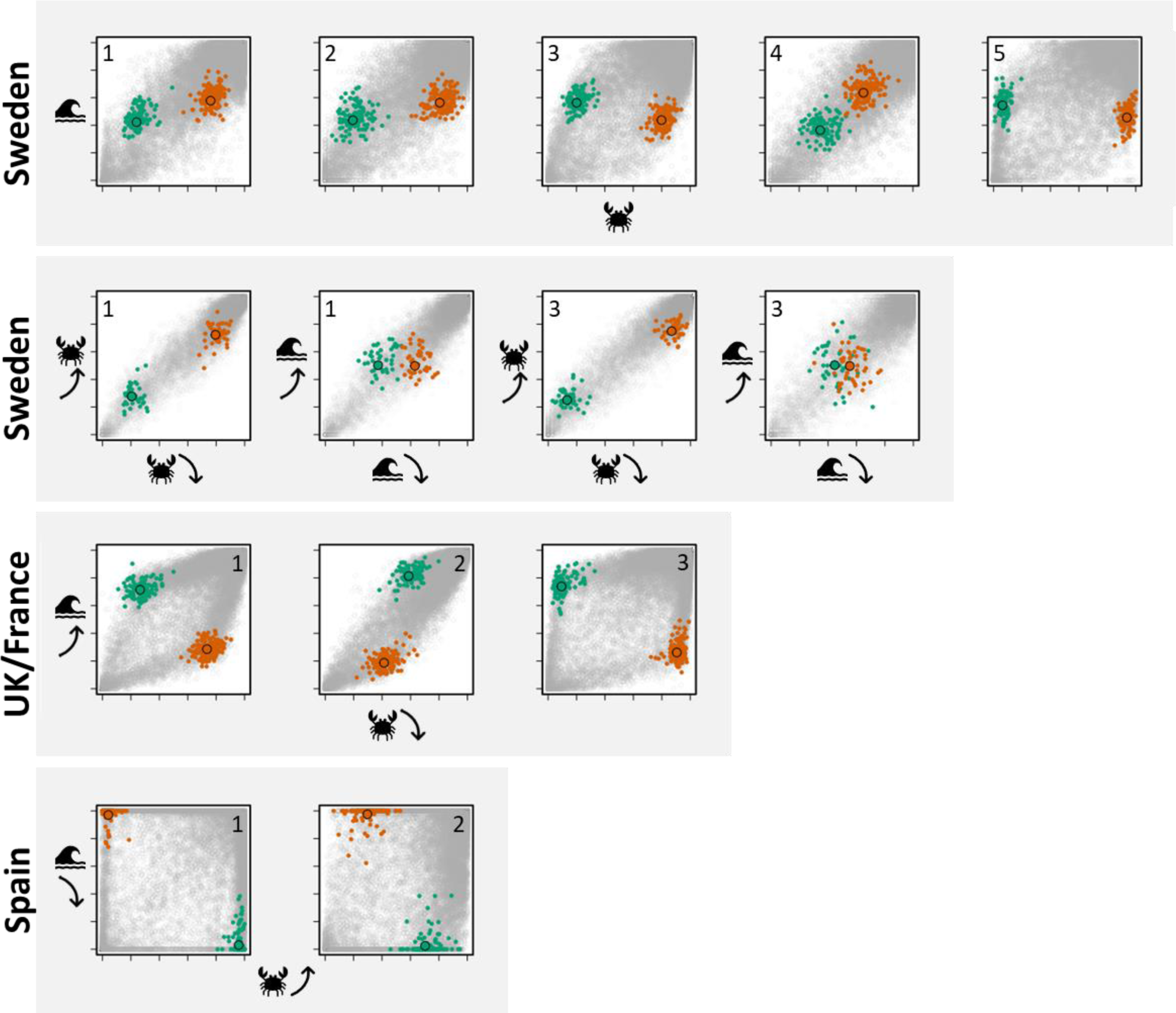
Allele frequency plots for the inversion region LGC9.1 for 14 different pairs of samples (axes ranging from 0 to 1; axis symbols as in Fig. 2). Each small point represents the frequencies of the reference genome allele for a SNP in the two local samples. Colored points represent arrangement-diagnostic SNPs (green: arrangement A; orange: arrangement B). Larger points with black borders indicate the frequencies of arrangement A and B obtained by averaging frequencies across arrangement-diagnostic SNPs. The geographical area of each sample pair is indicated on the left; the location ID is shown within each plot. Analogous plots for all other inversion regions can be found in Fig. S3.

After randomization, one of us scored 4cM windows along chromosomes by eye for the presence of a parallelogram. We then asked whether the regions with visible parallelograms correspond to regions containing known inversions (Westram et al. 2021). We found that many windows could not be scored visually (56% of windows were labelled “uncertain”), due to noisy data and / or small numbers of SNPs. However, where scoring was possible, inversions could be detected relatively reliably: Of 243 windows overlapping with an inversion that could be scored, 197 contained a detectable parallelogram (false negative rate 19%). Where no parallelogram was visible, this was often because arrangement frequency did not differ much between the two populations (as inferred below), and thus no parallelogram was expected despite the presence of an inversion polymorphism (Fig. 1D, Fig. S2). The false positive rate (parallelogram inferred without the presence of a known inversion) was 11% (42 out of 384 windows). For details, see Fig. S2.

### Estimation of arrangement frequencies

We determined a set of diagnostic SNPs for each arrangement. The numbers of such SNPs varied between twelve and 265 per inversion. By averaging the frequencies of diagnostic alleles, we estimated the frequencies of the A and B arrangement in each of the samples (Fig. 3 and Fig. S3). The frequencies obtained in this way generally corresponded very well to the visible parallelogram corners (Fig. 3 and Fig. S3) and were also very similar to frequencies obtained from individual sequencing data (Westram et al. 2021) (Fig. S4, Fig. S5), validating our pool-seq approach.

Note that the data for the Swedish Low-High pairs had been processed separately from the rest of the dataset and does not contain all diagnostic SNPs. Arrangement frequency estimates are therefore based on a reduced number of SNPs (Fig. S3). However, there were always at least 10 diagnostic SNPs, except for the frequencies of LGC6.1/2 and LGC10.1 (5 SNPs and 1 SNP, respectively), which therefore need to be treated with caution.

### Association of arrangements with selective axes

Our results show that all inversions known from studies in Sweden are also segregating in other European locations (Fig. 4). Inversions were almost always segregating at the location level, i.e., usually both arrangements were present in each location, with the rarer arrangement having a frequency higher than 0.05. A notable exception is LGC17.1, which was near fixation overall in UK/France (Fig. 4). In contrast, within the Crab or Wave ecotype within a given location, near-fixation was more common. In particular, the two Spanish Wave ecotype samples were close to fixation for one arrangement for most inversions (Fig. 4).

**Fig. 4:**
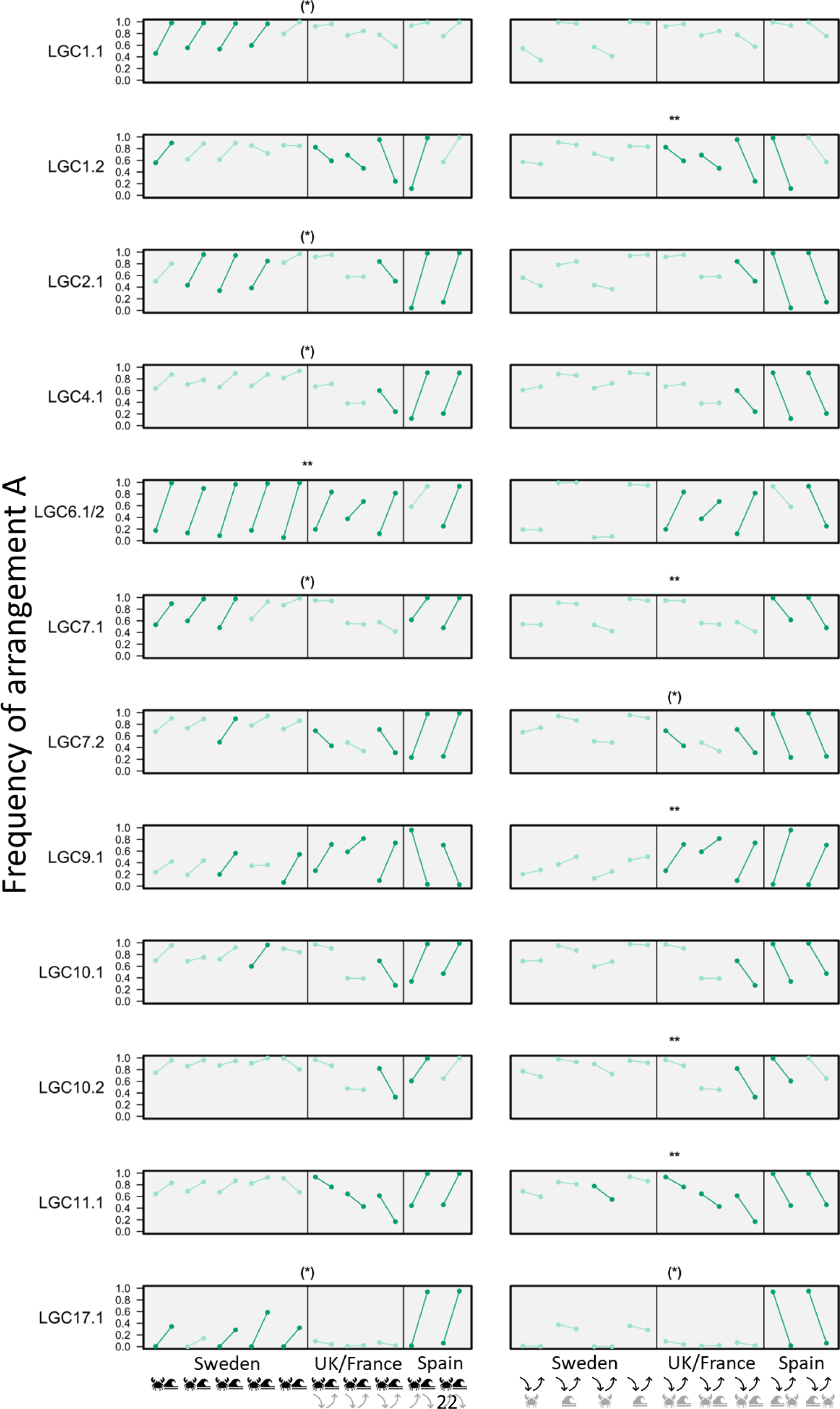
Frequency of arrangement A for each inversion, location, and sample. Lines connect samples from the same location. Samples within locations are shown in the order Crab-Wave (left) or Low-High (right). Inversions associated with Crab-Wave (Low-High) are expected to show consistent directionalities of the lines in the left (right) plot. Dark green color indicates significantly elevated arrangement frequency difference between environments within locations. Asterisks indicate the significance of a Wilcoxon signed rank test for general frequency differences between Crab and Wave / Low and High (* p<0.05; ** p<0.01; *** p<0.001; brackets: not significant after sequential Bonferroni correction).

All inversions showed elevated differentiation (when compared to collinear SNPs) between samples in at least three locations (Fig. 4, dark green lines). Within geographical areas (Sweden, France/UK, Spain), the direction of differentiation was always consistent (when considering only locations showing elevated differentiation according to our test), e.g., the Crab sample always had a higher frequency of arrangement A than the Wave sample of the same location.

We compared patterns between geographical areas to reveal whether arrangements were consistently associated with the Crab-Wave axis or the Low-High axis. Wilcoxon signed rank tests indicated that out of 12 inversions analyzed, 11 showed significant consistent associations with at least one of the two environmental axes (six remained significant after sequential Bonferroni correction; Fig. 4). Four inversions were significantly associated with the Crab-Wave axis, 5 with the Low-High axis, and two with both axes. However, it is clear that in some of these cases, while directionalities are similar across countries, local arrangement frequency differences between environments are often small and not indicative of selection (light green lines in Fig. 4; e.g. LGC7.1 Low-High). Clear environment-inversion associations consistent across countries with at least 5 instances of elevated differentiation within locations (dark green lines in Fig. 4) were found in five cases: LGC6.1/2 (Crab-Wave), LGC7.1 (Crab-Wave), LGC17.1 (Crab-Wave), LGC9.1 (Low-High), and LGC11.1 (Low-High).

### Analysis of arrangement content

We analyzed the arrangement content for two inversions that showed particularly strong associations with an environmental axis, LGC6.1/2 (associated with Crab-Wave) and LGC9.1 (associated with Low-High). We found that most SNPs associated with the inversion in one geographical area also showed a similar association in the other two geographical areas (Fig. 5, Fig. S6). This is consistent with the fact that for all inversions we could find “globally” diagnostic SNPs that work across all studied countries. However, we also found some SNPs that were associated with the inversion in one area but invariant in other areas (lines in grayish colors in Fig. 5, Fig. S6). In addition, some SNPs showed reverse associations with the inversion in different countries, particularly when Spain was compared to UK/France and Sweden (red lines in Fig. 5, Fig. S6).

**Fig. 5:**
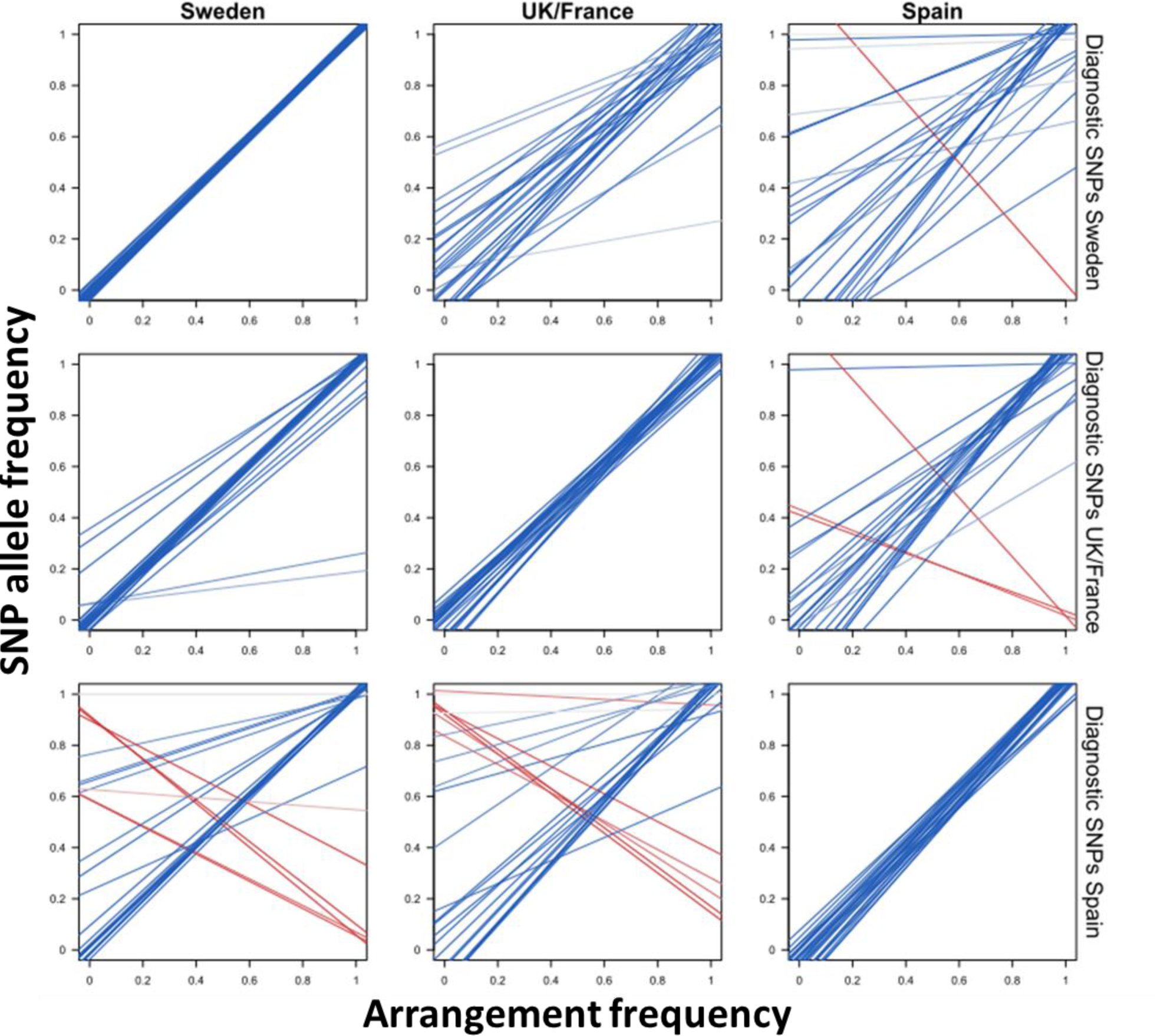
Patterns of locally arrangement-diagnostic SNPs in other geographical areas for the LGC6.1/2 inversion. Each line represents the regression of SNP allele frequency against the frequency of arrangement A for one SNP. The geographical area where the SNPs are arrangement-diagnostic is indicated on the right (SNPs can be arrangement-diagnostic in multiple countries, so they can occur in multiple rows); regression lines always show the allele that is positively associated with arrangement A in this geographical area. Line color indicates Pearson’s correlation coefficient (r), with a color gradient from negative (red) via 0 (grey) to positive (blue).

## Discussion

### Using SNP allele frequency data to analyze inversion regions

The importance of chromosomal rearrangements in adaptive divergence has become increasingly clear, and testing for their presence and frequency in natural populations is important for understanding adaptive processes. In particular, it is a largely open question whether inversions contribute to adaptation repeatedly across large geographical scales, and whether they differ in adaptive content between geographical areas, representing versatile “toolboxes”. While multiple methods exist to determine the presence and population frequency of inversions, they usually require individual sequencing data (which can generate high costs if many locations are studied) and sophisticated analyses. Here, we present ideas of how simple allele frequency data (here, cost-effective pool-seq data) could be used to study inversions.

Our approach relies on a parallelogram shape in plots of allele frequencies in population 2 against population 1, which appears when populations are connected by gene flow but differ in arrangement frequency for an inversion. These parallelograms are essentially an outcome of the extensive LD in the inversion region, generating associated frequency shifts across many SNPs. Our simulations and empirical results thus demonstrate that LD information is not entirely lost in allele frequency data, as sometimes assumed, and extends beyond the read- or paired read-level previously targeted by LD analyses using pool-seq data (Feder et al. 2012). The parallelograms are useful both for de-novo detection (presence of a parallelogram indicates presence of a polymorphic inversion) and frequency determination (parallelogram corners indicate arrangement frequencies) and thus provide an alternative to often more costly and labor-intensive methods based on cytogenetics, LD analysis or long-read sequencing data.

Inversion de-novo detection by eye worked best for sample pairs with limited general differentiation, many individuals per pool, and larger arrangement frequency differences between populations. LD-based approaches in individual data are certainly generally more reliable and would often be required to confirm findings from pool-seq data. However, we note that for high-quality samples parallelogram patterns are very clear and many rearrangements can be discovered with little doubt. For example, our French pools showed very little noise and all inversions were reflected by extremely clear parallelograms that not only allowed for accurate inversion detection but also for reliable frequency estimation (Fig. S3). We conclude that for researchers studying local adaptation using pool-seq data, it is certainly worth screening the genome for inversions using allele frequency plots. Regions with clear parallelograms will usually also show elevated F_ST_; however, the advantage of the parallelogram method is that a parallelogram clearly hints at a genomic rearrangement (while high F_ST_ can also occur in collinear regions), and in contrast to F_ST_ it allows for arrangement frequency estimation. Allele frequency plots could also be a quick way to scan a genome for candidate inversion regions even if individual sequencing data is available, being faster than a full LD analysis. One important thing to note here is that other chromosomal rearrangements, e.g., large duplications, might potentially also lead to parallelogram shapes. Therefore, studies relying on de-novo detection (in contrast to our study, where inversions were previously known), should check the type of rearrangement in follow-up analyses, potentially using individual sequencing data.

Determining the location of parallelogram corners visually, or using allele frequencies of diagnostic SNPs, it is possible to estimate arrangement frequencies in each sample. Applying the latter approach, we obtained arrangement frequency estimates that were highly reliable. To find “arrangement-diagnostic SNPs”, we made use of the fact that we had sampled multiple population pairs, identifying the SNPs most strongly associated with parallelogram corners across different locations. Using these diagnostic SNPs, it was possible to detect inversion polymorphism even in locations with very noisy parallelograms (e.g., Fig. 3, Sweden 1 Crab-Wave) and in locations where arrangement frequencies were very similar in the two samples and thus no parallelogram appeared (e.g., Fig. 3, Sweden 4). Finding arrangement-diagnostic SNPs directly from pool-seq data thus provides an alternative to first determining them from individual sequencing data (Kapun et al. 2014). Of course, even SNPs detected in this way might not be fully reliable – for example, SNPs that are arrangement-diagnostic in 9 locations might not be so it the 10^th^ location, and while our approach finds statistically associated SNPs, unless very stringent thresholds are used this association may not be perfect (see SNPs deviating from the parallelogram corners in Fig. S3). Nevertheless, if multiple putatively arrangement-diagnostic SNPs are used, reliable estimates should be possible.

It is likely that some SNPs violate the assumptions of being variable in one arrangement only and not strongly differing in within-arrangement frequency between local populations (see Rationale). However, these violations only introduce noise; as arrangement frequency estimation relies on SNPs found consistently near the parallelogram corners, these SNPs do not strongly affect our approach. Similarly, some collinear SNPs accidentally included when inversion boundaries are not exactly known, or because of assembly errors, may generate points inside the parallelogram but should not generate major problems in frequency estimation. However, inversions affected by very large amounts of gene conversion or double cross-over might be harder to analyze as this will generate a larger number of SNPs “inside” the parallelogram.

In this paper, we have aimed at introducing the general rationale of using allele frequency data for inversion frequency estimation and demonstrating its usefulness in an empirical system. We have not developed a fully automated approach to detect and quantify inversion polymorphism, but rather use a heuristic approach. Automation could be an interesting possibility for the future. A challenge for such a tool is the incorporation of SNPs violating assumptions (see above) as well as sampling noise; in our pilot attempts, automation (results not shown) always performed much worse than the heuristic approach presented here and even just visual examination.

Extensive simulations exploring the limits of parallelogram detection (visually or via a heuristic approach) and optimizing parameter settings for the detection of diagnostic SNPs are well beyond the scope of this paper. However, they could be an interesting avenue for future work. Relevant factors to vary include the extent of gene flow between populations, the inversion histories (e.g., time to accumulate differences), the level of gene flux between arrangements, and the extent of noise in the allele frequency data (e.g., numbers of individuals in the pool; relative DNA contribution of each individual to the pool).

### Inversions across large geographical scales in *Littorina*

In our study system, we used the approach discussed above to confirm the presence of polymorphic inversions across Europe, determine arrangement frequencies (which was impossible based on earlier FST approaches; Morales et al. 2019) and analyze arrangement content.

A first main finding is that the known inversions are polymorphic across most of the studied European locations. This extends the work on inversion polymorphism in Sweden (Faria et al. 2019a; Westram et al. 2021) and earlier pool-seq work, which could often show high divergence in inversion regions, but could not detect inversion polymorphism in locations where the different ecotypes did not differ in arrangement frequency (e.g. LGC7.1 in the UK and France). This widespread polymorphism might indicate balancing selection on at least some of the inversions (Wellenreuther and Bernatchez 2018; Faria et al. 2019a, 2019b).

One obvious question is whether parallel evolution of Crab-Wave and Low-High ecotypes in *Littorina* is driven by the same inversions across Europe. Using the arrangement frequency data in this study we could address this question. We found that 5 arrangements are clearly associated with either the Crab-Wave (LGC6.1/2, LGC7.1, LGC17.1) or the Low-High (LGC9.1, LGC11.1) selection axis, while other inversions show less clear associations. These results are consistent with work on other systems showing the same arrangement repeatedly associated with the same environment in different geographical locations (e.g. Mérot et al. 2018; Christmas et al. 2019), and also overall consistent with earlier analyses of the same dataset (Morales et al. 2019). However, this previous work did not infer arrangement frequencies and relied on elevated F_ST_ in inversion regions. F_ST_ in inversion regions is affected by various factors, including not only arrangement frequency but also inversion age, selection, and the extent of gene flux. Here, we obtain arrangement frequency estimates that are independent of these factors and therefore more reliable (Fig. 4). However, the associations with environmental axes still need to be interpreted with some caution: It is not possible to fully disentangle the Crab-Wave and the Low-High selective axes, partly because locations within geographical areas all have the same association between environmental axes (Fig. 2) but are treated as independent here. It is only possible to resolve this with a modified sampling design, where Crab and Wave individuals are sampled at exactly the same shore level, and Low and High individuals are sampled within Wave or Crab, for each of the geographical area.

Arrangement-environment associations that are consistent between geographical areas might partly be driven by a set of selected SNPs strongly differentiating the arrangements across different locations. Finding causal SNPs under selection in high-LD regions like inversions is notoriously difficult as many SNPs show correlated patterns (Ayala et al. 2017). However, as part of our approach to determine arrangement frequencies, we determined “ arrangement-diagnostic” SNPs associated with arrangements across multiple locations; these are the strongest candidates for SNPs driving consistent arrangement frequency differences and warrant further study. Thus, comparisons across multiple geographical locations might be one way of narrowing down candidate SNPs within inversion regions and our approach makes this feasible without the high cost of individual sequencing.

Interestingly, most inversions studied here, even if significantly associated with a single selective axis, did not show fully consistent associations across all locations. For example, LGC7.2 was significantly associated with the Low-High axis, but visual inspection also suggests a Crab-Wave association pattern in some Swedish locations. Associations with multiple selective axes were already indicated in Morales et al. (2019). For LGC17.1, the clear association with the Crab-Wave axis is limited to Sweden and Spain but absent in the UK/France. These examples of differences between geographical areas might indicate location-specific patterns or strengths of selection. We use a crude categorization of samples into “Crab” and “Wave” or “Low” and “High” habitats, but the details and strengths of selective pressures likely vary between countries. In stickleback lake-stream divergence, Stuart et al. (2017) discovered that non-parallel divergence patterns can be attributed to slight variations in local selection pressures. These nuances are often overlooked when habitats are simply categorized as “lakes” and “streams”. Such effects could explain some of the complex patterns we see.

Additionally, or alternatively, the differences between geographical areas could indicate that inversions have different functional contents in different locations – i.e., contain different “tools” adjusted to local requirements. For example, LGC9.1 might contain only Low-High adaptive variants in UK/France and Spain but might additionally contain Crab-Wave adaptive variants in Sweden. In line with this, we find that some of the SNPs differentiating arrangements in Sweden do not show such associations in the other geographical areas (Fig. S6).

Such differences in arrangement content were generally found when focusing on locally arrangement-diagnostic SNPs in LGC6.1/2 and LGC9.1. Each of the geographical areas has a small number of SNPs that do not differentiate arrangements elsewhere. These could contribute to location-specific adaptive differences (e.g., differences in shell sculpture that exist in Spain but not elsewhere), contribute to adaptation to environmental contrasts that do not exist in the other areas, or fulfil adaptive functions that are fulfilled by other parts of the genome elsewhere. While we cannot currently say whether any of these SNPs are actually functional, these results nevertheless demonstrate the potential for geographical variation in arrangement content. In particular, we find some SNPs with reversed associations with the inversion in the UK/France vs. Spain for both of the studied inversions. In LGC6.1/2 (which is generally associated with the Crab-Wave axis), the genomic regions around these SNPs could contribute to adaptation to the Low-High axis, which is reversed between the UK/France and Spain. Similarly, in LGC9.1 (which is generally associated with the Low-High axis), these regions could contribute to Crab-Wave adaptation.

These results and hypotheses emphasize the need to analyze patterns of selection within arrangements. While some studies have focused on selection within arrangements – for example, an experimental study on *Drosophila subobscura* showed how markers within arrangements changed more than expected by chance in the course of a selection experiment (Santos et al. 2016) – arrangement content remains largely unexplored for many systems with inversion-facilitated adaptation.

In summary, we show that allele frequency data, e.g., from pool-seq, has the potential to increase our understanding of the role of inversions in adaptive evolution. Allele frequency plots are straightforward to generate, by-eye analysis is sufficient in some cases, and heuristic approaches are easy to implement. These plots can be used to detect undiscovered inversions, determine arrangement frequencies, and analyze inversion content. Applying these ideas to the *L. saxatilis* system, we obtain results strongly consistent with inversions being associated with divergent adaptation, as predicted by theory and suggested by previous work on this system (Morales et al. 2019). We find that while arrangements clearly show repeated associations with the same environmental axes, there are also differences between geographical areas and variation in arrangement content. This highlights the need to go beyond focusing on an inversion as a single unit: While inversions might be useful toolboxes for adaptation, what matters are the tools inside.

## Supporting information

All Supplementary Material

## Data and Code Accessibility Statement

Raw sequencing reads can be found in the Sequence Read Archive (BioProject PRJNA494650). The allele count data from Morales et al. (2019), used as input in this study, can be found on Dryad (doi…). R scripts for all customs steps in this manuscript are deposited in GitHub (link…).

