## Supplementary material for "Understanding the adaptive role of chromosomal inversions across large geographical scales: The potential of pool-seq data": All Supplementary Material

### *Supplementary Figures*

**LG1 / LGC1.1:**

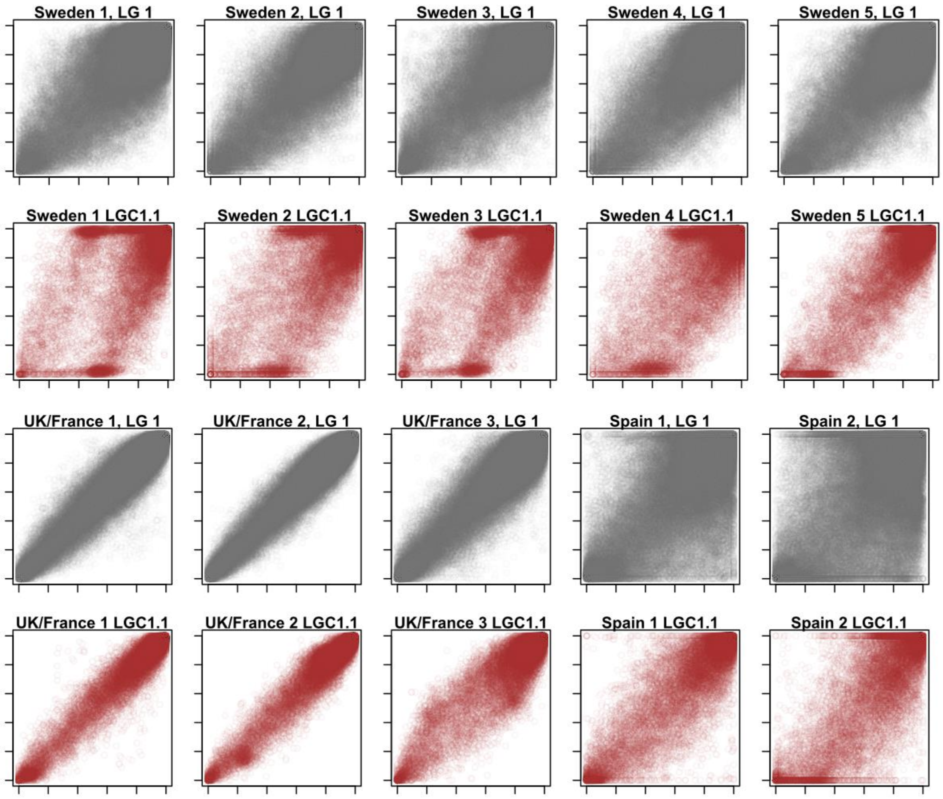


**LG1 / LGC1.2:**

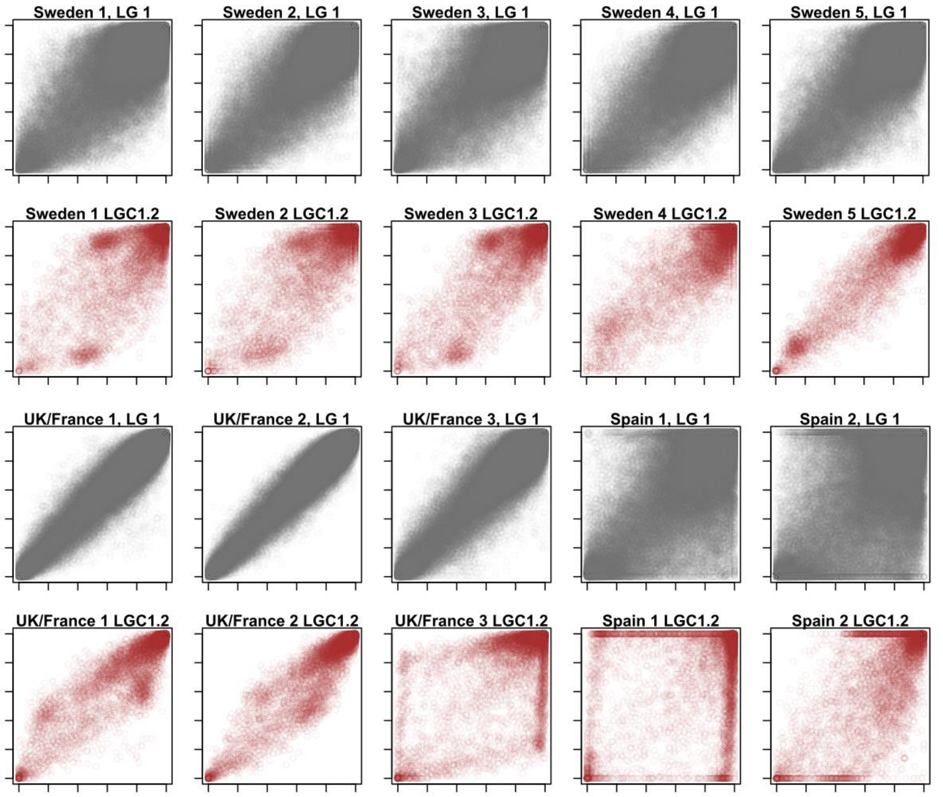

#### LG2 / LGC2.1:

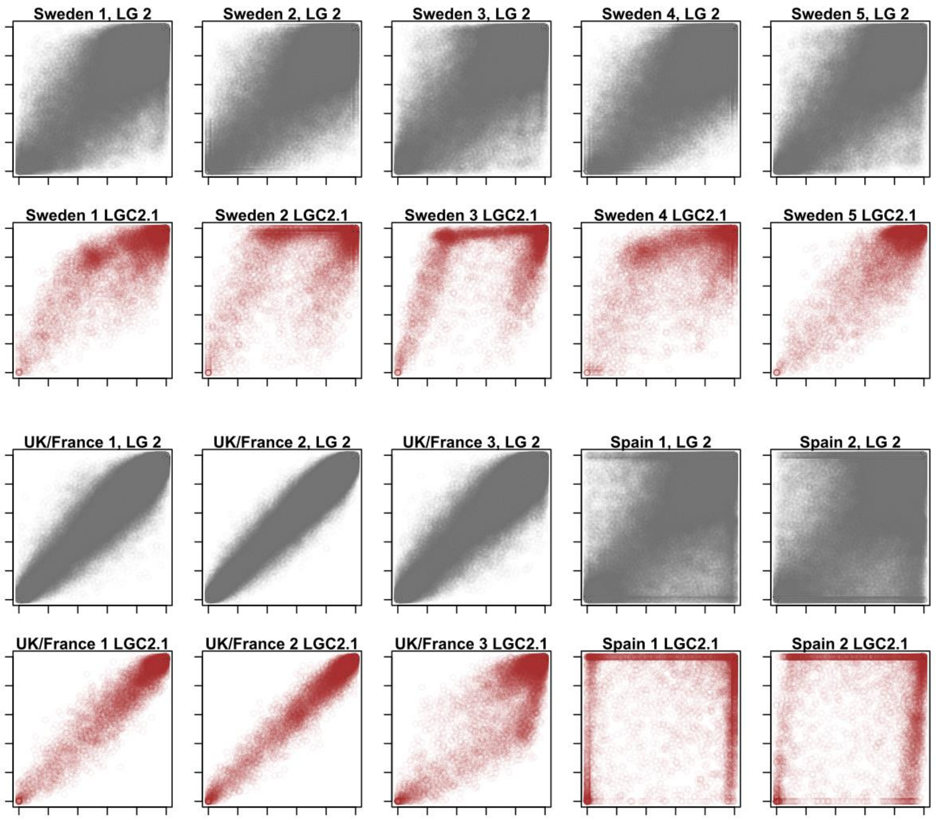


#### LG4 / LGC4.1:

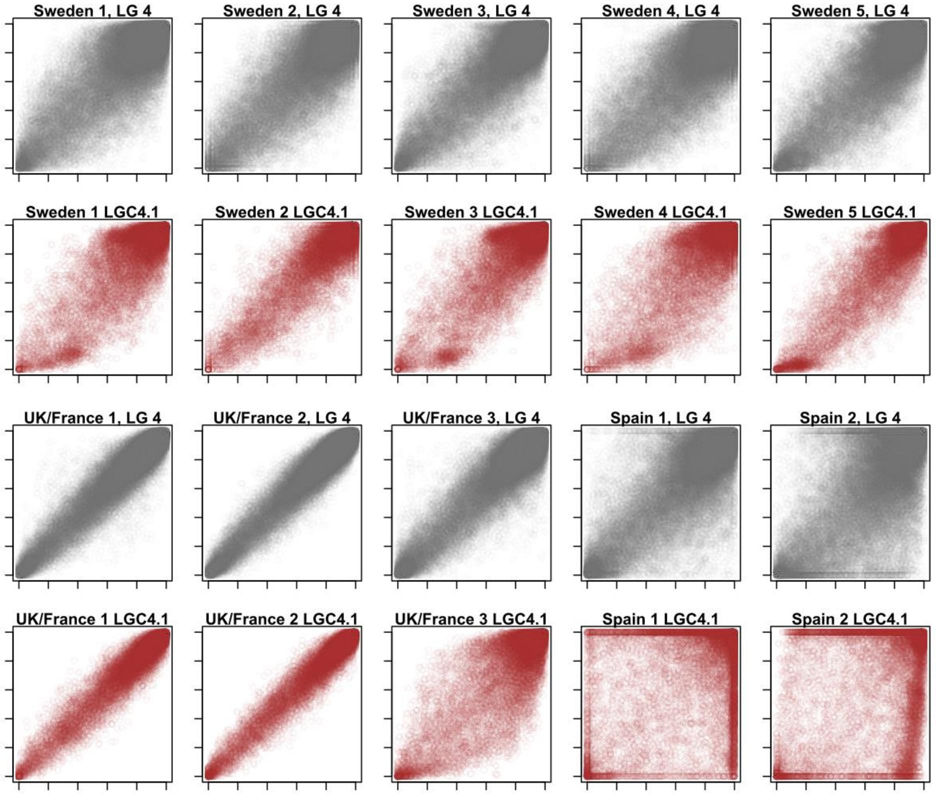

**LG6 / LGC6.1/2:**

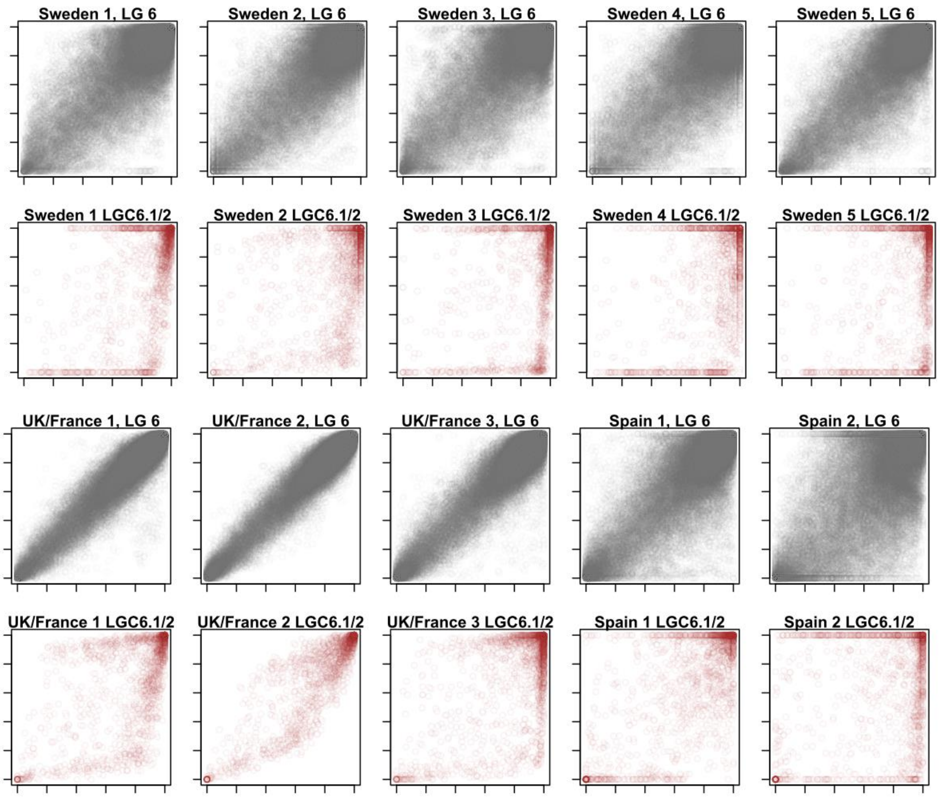


**LG7 / LGC7.1:**

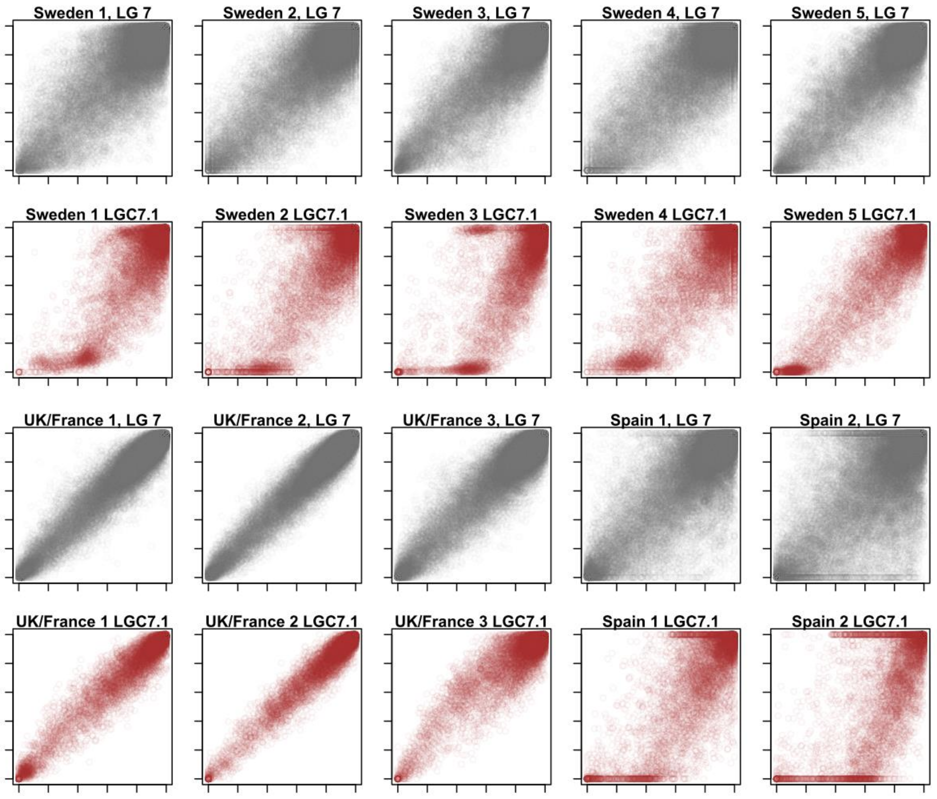

#### LG7 / LGC7.2:

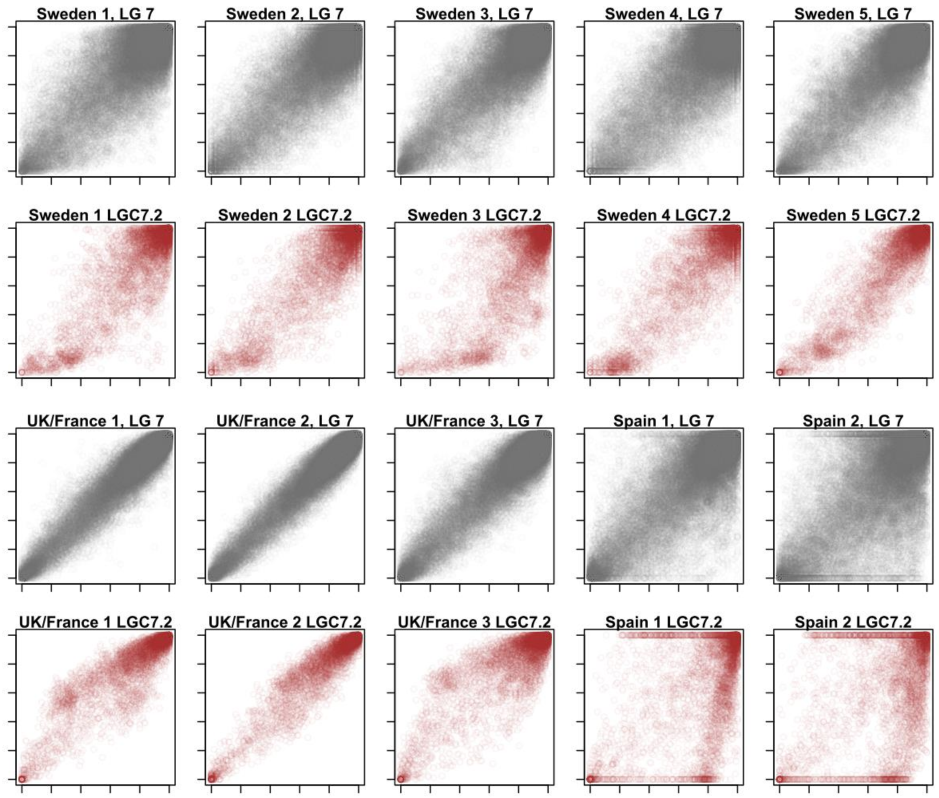


#### LG9 / LGC9.1:

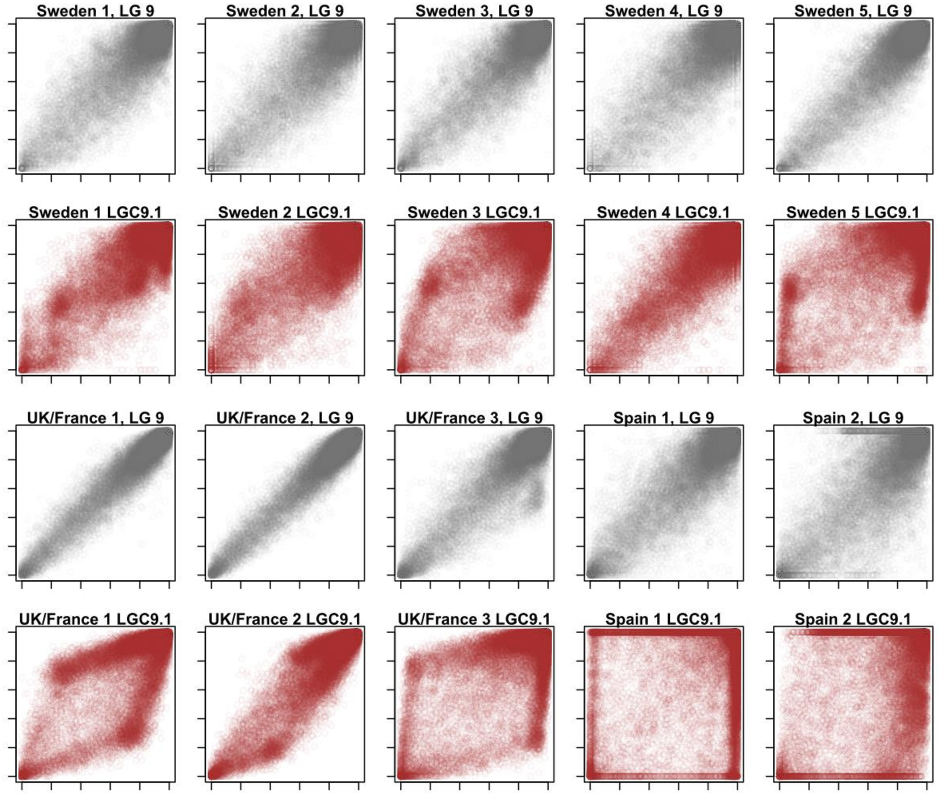

**LG10 / LGC10.1:**

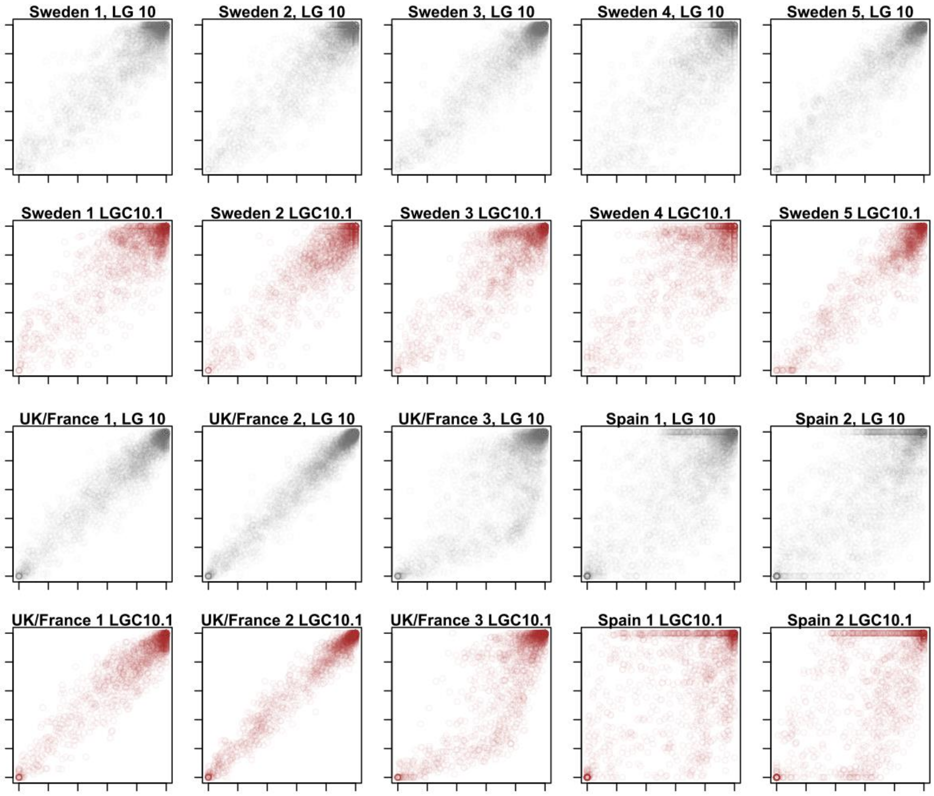


**LG10 / LGC10.2:**

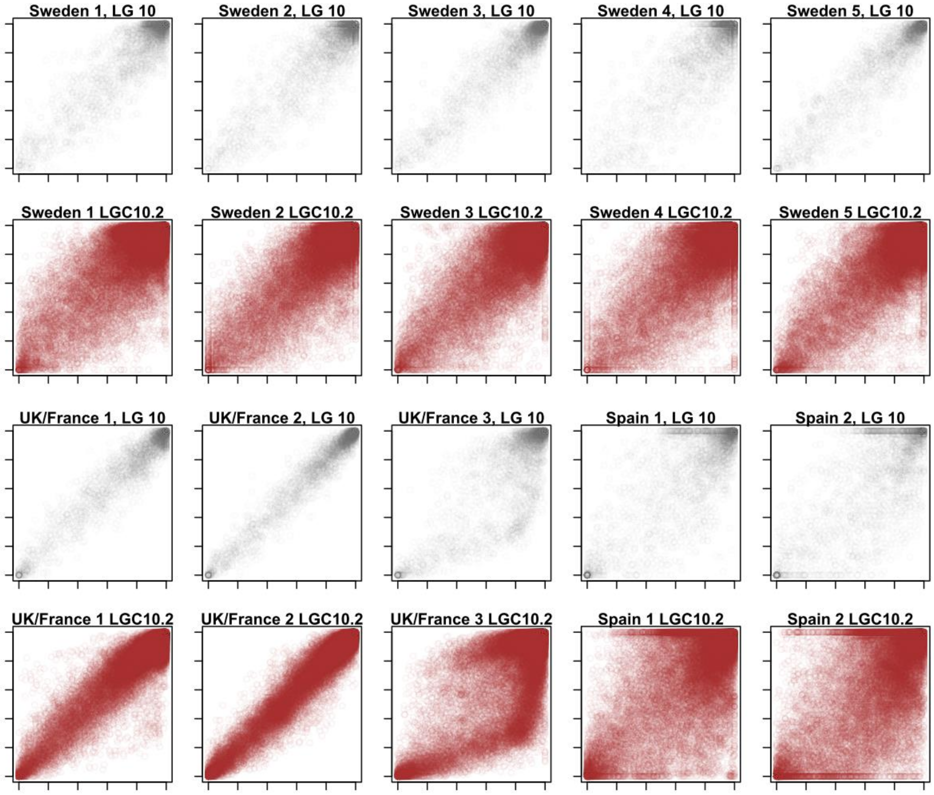

**LG11 / LGC11.1:**

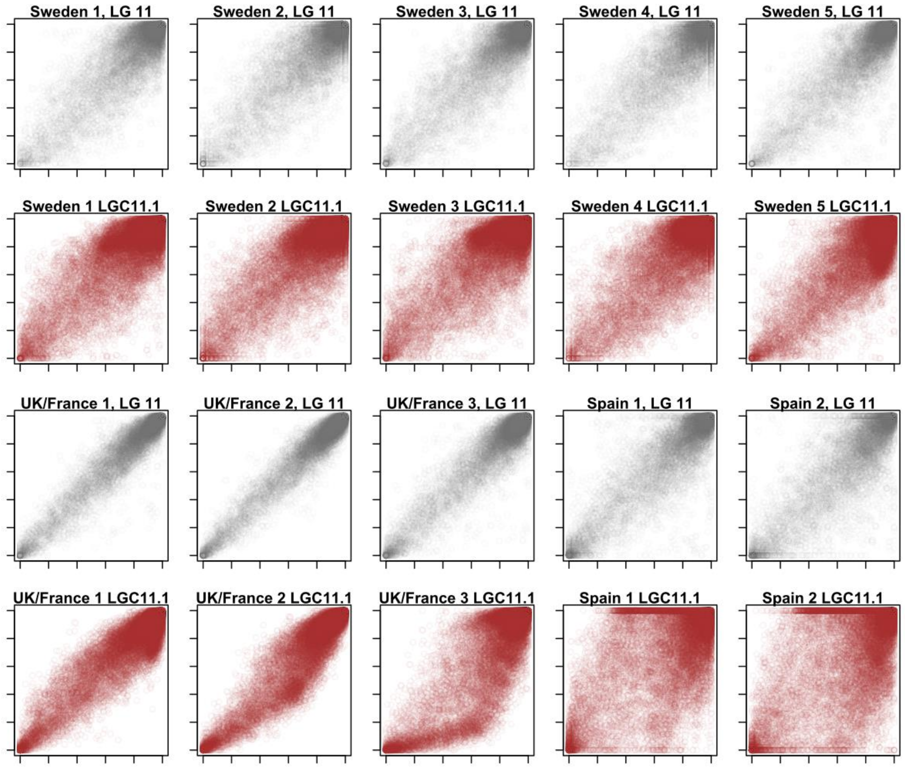


**LG17 / LGC17.1:**

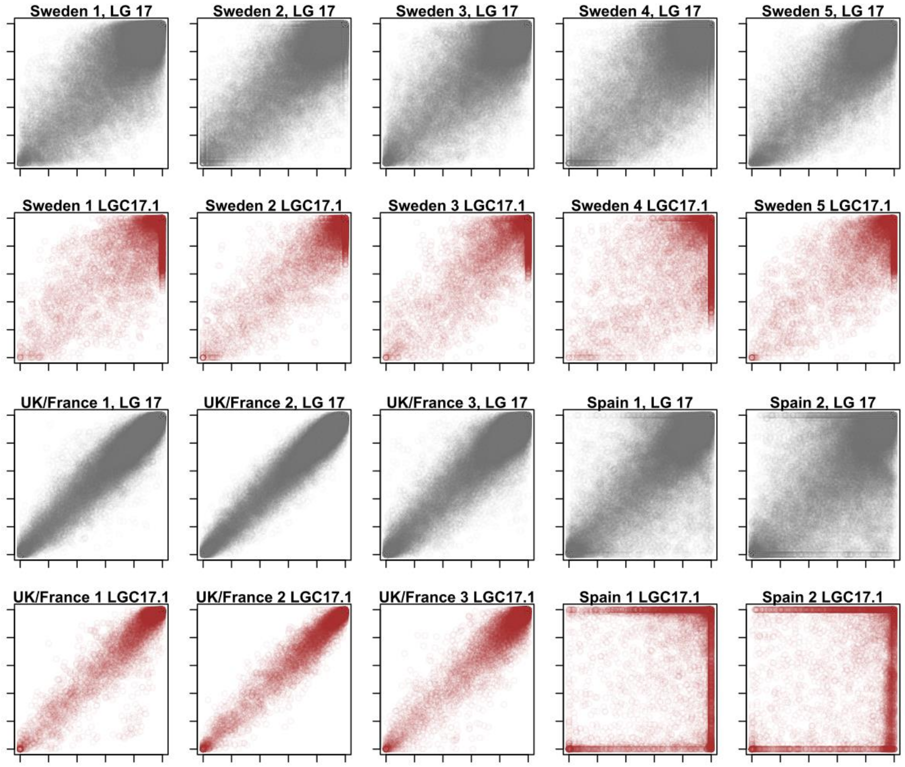


Fig. S1: All Crab-Wave allele frequency plots for known inversion regions and collinear regions on the same linkage group. Each plot shows the Wave allele frequency against the Crab allele frequency for all SNPs in the collinear region of the linkage group (grey) or all SNPs in the focal inversion region (red), with axes going from 0 to 1. Linkage group / inversion IDs are given above each plot.

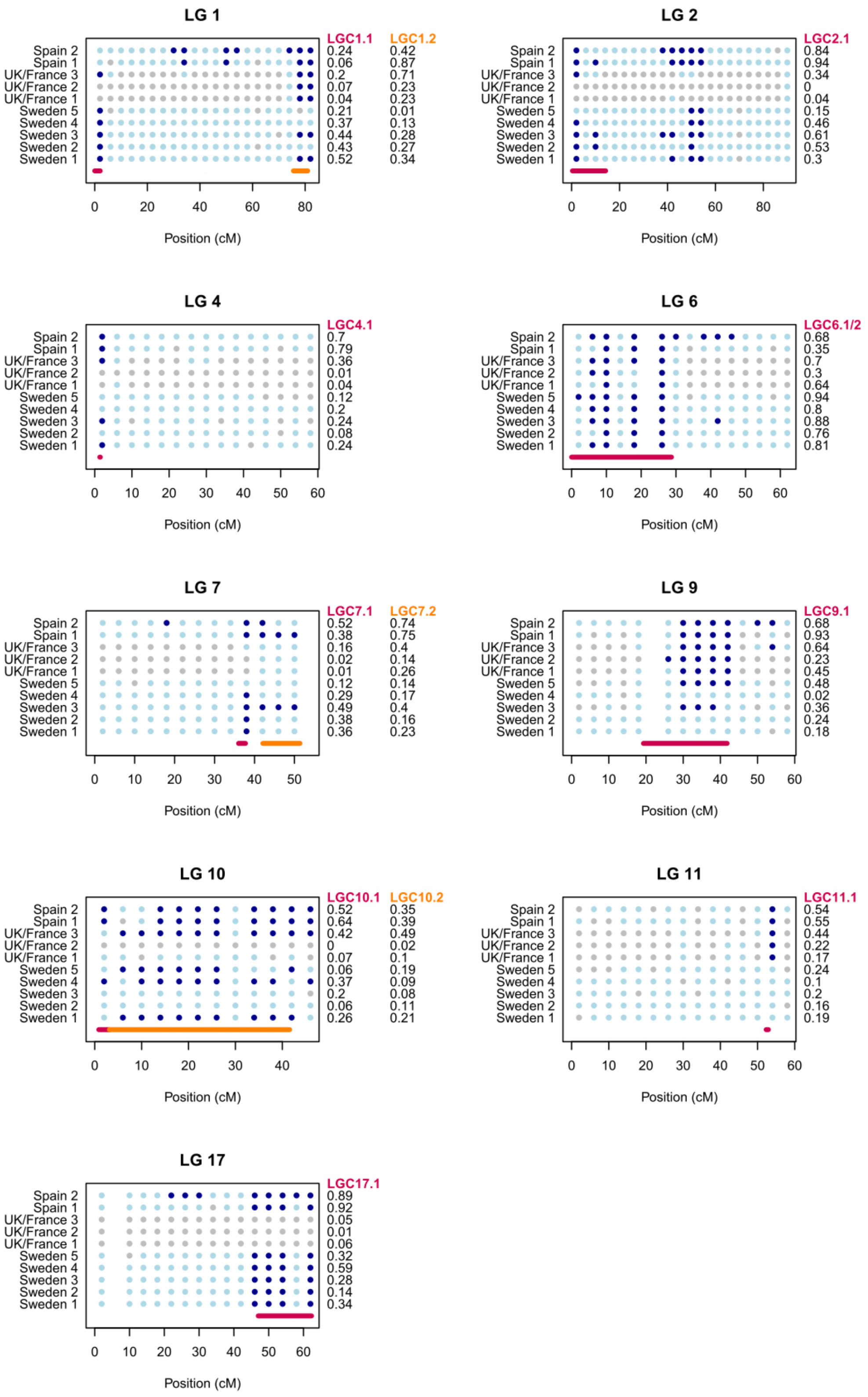


Fig. S2: Results of inversion de-novo detection for the nine linkage groups containing known inversions. Crab-Wave allele frequency plots were scored for parallelograms by eye in 4cM genomic windows. Each point reflects one such genomic window for a given location. Dark blue points indicate a visible parallelogram, grey points no visible parallelogram, and light blue points uncertainty. Gaps indicate missing data. Horizontal lines highlight the true positions of the inversions, based on Westram et al. (2021). The numbers at the right side of the plots reflect the absolute arrangement frequency difference between Crab and Wave for each location and inversion. These numbers demonstrate that inversions are often missed when the arrangement frequency difference is small, so that no clear parallelogram is expected.

#### LGC1.1:

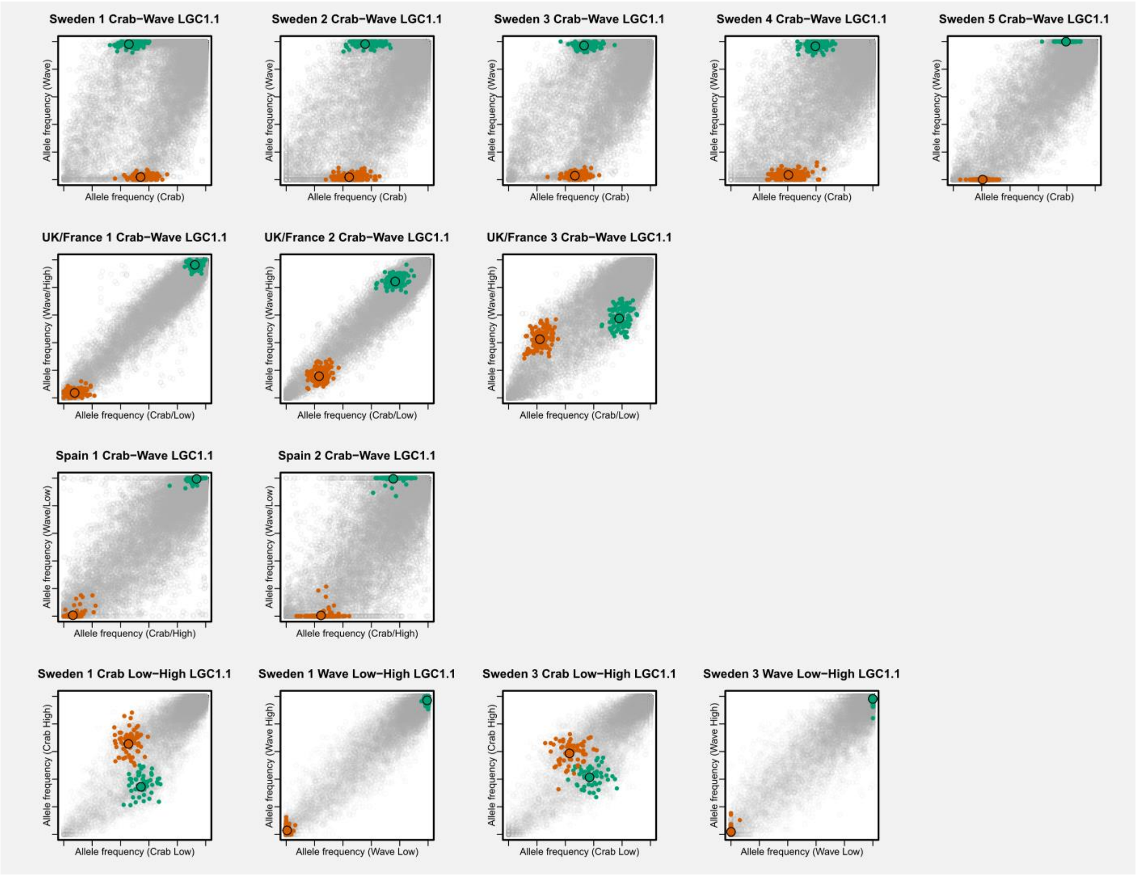

Sweden 1 Crab−Wave LGC1.1
Sweden 2 Crab−Wave LGC1.1
Sweden 3 Crab−Wave LGC1.1
Sweden 4 Crab−Wave LGC1.1
Sweden 5 Crab−Wave LGC1.1
Allele frequency (Wave)
Allele frequency (Crab)
UK/France 1 Crab−Wave LGC1.1
UK/France 2 Crab−Wave LGC1.1
UK/France 3 Crab−Wave LGC1.1
Allele frequency (Wave/High)
Allele frequency (Crab/Low)
Spain 1 Crab−Wave LGC1.1
Spain 2 Crab−Wave LGC1.1
Allele frequency (Wave/Low)
Allele frequency (Crab/High)
Sweden 1 Crab Low−High LGC1.1
Sweden 1 Wave Low−High LGC1.1
Sweden 3 Crab Low−High LGC1.1
Sweden 3 Wave Low−High LGC1.1
Allele frequency (Crab High)
Allele frequency (Crab Low)
Allele frequency (Wave High)
Allele frequency (Wave Low)


#### LGC1.2:

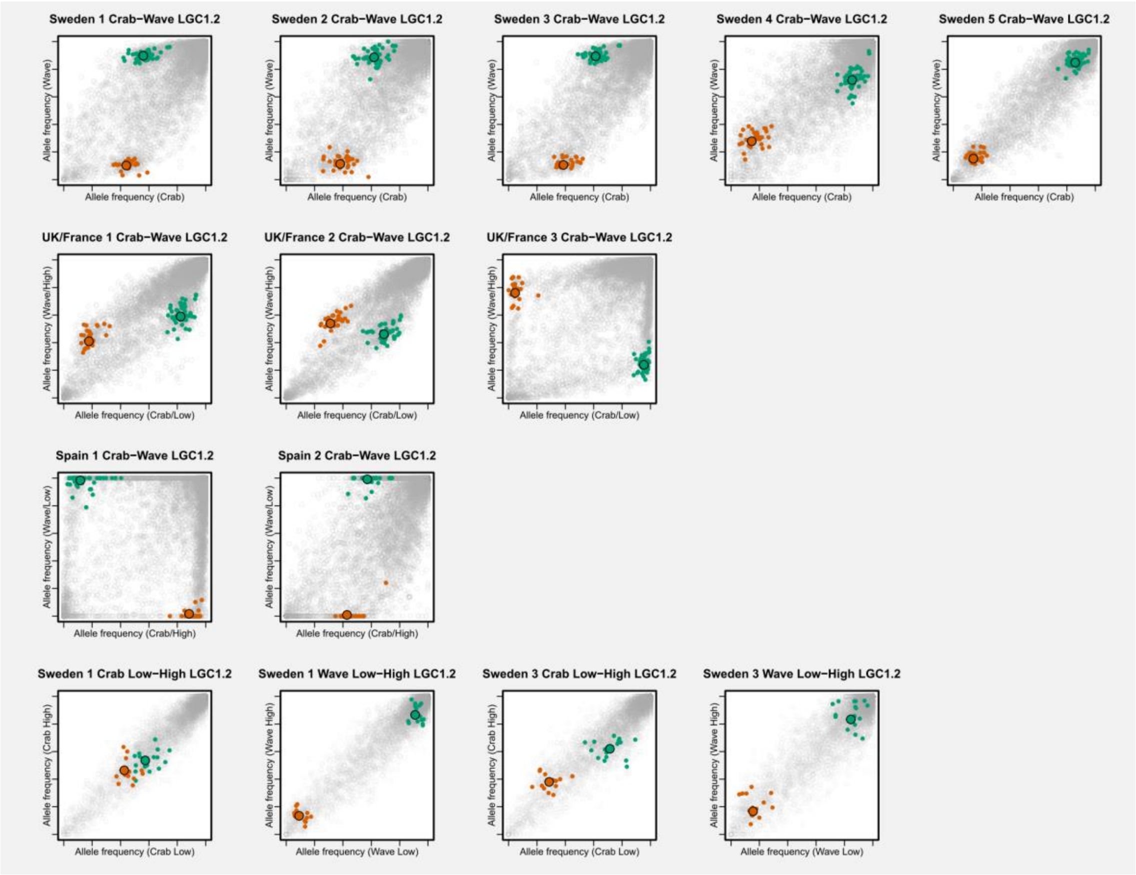

Sweden 1 Crab−Wave LGC1.2
Sweden 2 Crab−Wave LGC1.2
Sweden 3 Crab−Wave LGC1.2
Sweden 4 Crab−Wave LGC1.2
Sweden 5 Crab−Wave LGC1.2
Allele frequency (Wave)
Allele frequency (Crab)
UK/France 1 Crab−Wave LGC1.2
UK/France 2 Crab−Wave LGC1.2
UK/France 3 Crab−Wave LGC1.2
Allele frequency (Wave/High)
Allele frequency (Crab/Low)
Spain 1 Crab−Wave LGC1.2
Spain 2 Crab−Wave LGC1.2
Allele frequency (Wave/Low)
Allele frequency (Crab/High)
Sweden 1 Crab Low−High LGC1.2
Sweden 1 Wave Low−High LGC1.2
Sweden 3 Crab Low−High LGC1.2
Sweden 3 Wave Low−High LGC1.2
Allele frequency (Crab High)
Allele frequency (Crab Low)
Allele frequency (Wave High)
Allele frequency (Wave Low)

#### LGC2.1:

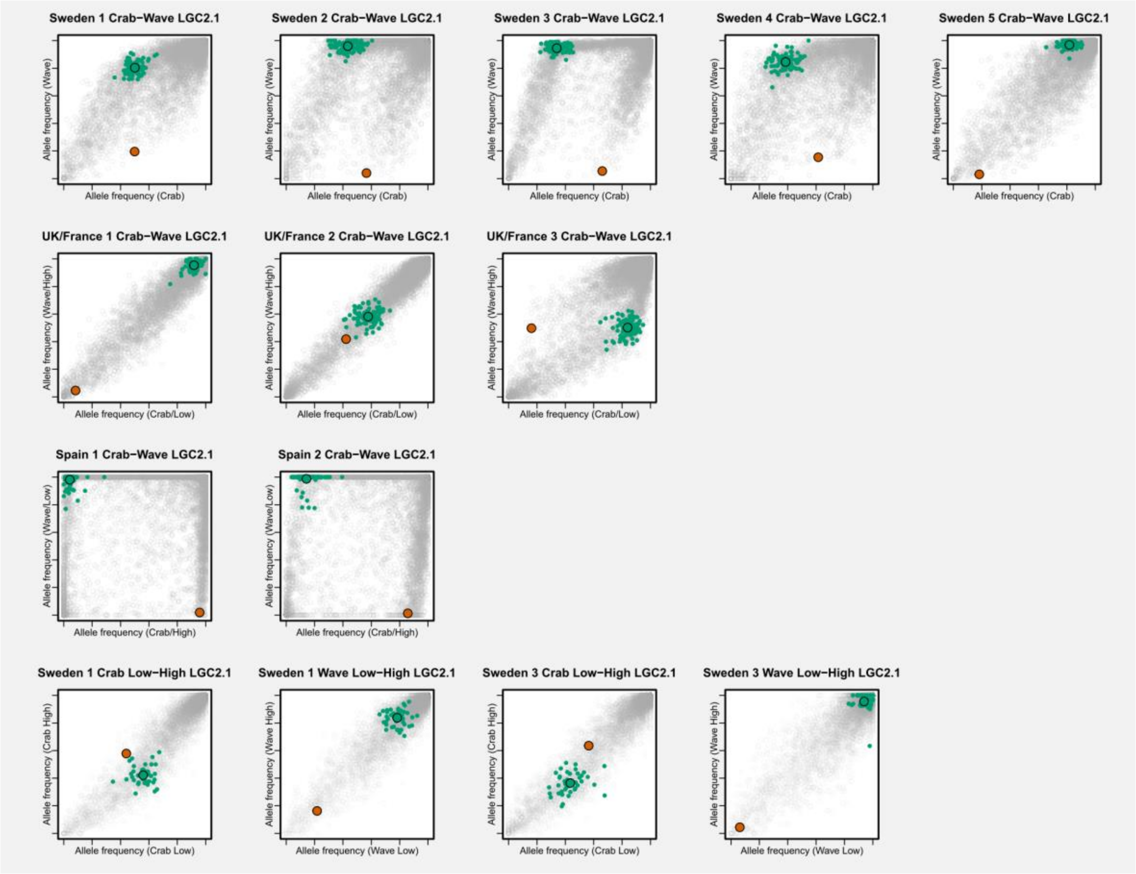

Sweden 1 Crab–Wave LGC2.1
Sweden 2 Crab–Wave LGC2.1
Sweden 3 Crab–Wave LGC2.1
Sweden 4 Crab–Wave LGC2.1
Sweden 5 Crab–Wave LGC2.1
Allele frequency (Wave)
Allele frequency (Crab)
UK/France 1 Crab–Wave LGC2.1
UK/France 2 Crab–Wave LGC2.1
UK/France 3 Crab–Wave LGC2.1
Allele frequency (Wave/High)
Allele frequency (Crab/Low)
Spain 1 Crab–Wave LGC2.1
Spain 2 Crab–Wave LGC2.1
Allele frequency (Wave/Low)
Allele frequency (Crab/High)
Sweden 1 Crab Low–High LGC2.1
Sweden 1 Wave Low–High LGC2.1
Sweden 3 Crab Low–High LGC2.1
Sweden 3 Wave Low–High LGC2.1
Allele frequency (Crab High)
Allele frequency (Crab Low)
Allele frequency (Wave High)
Allele frequency (Wave Low)


#### LGC4.1:

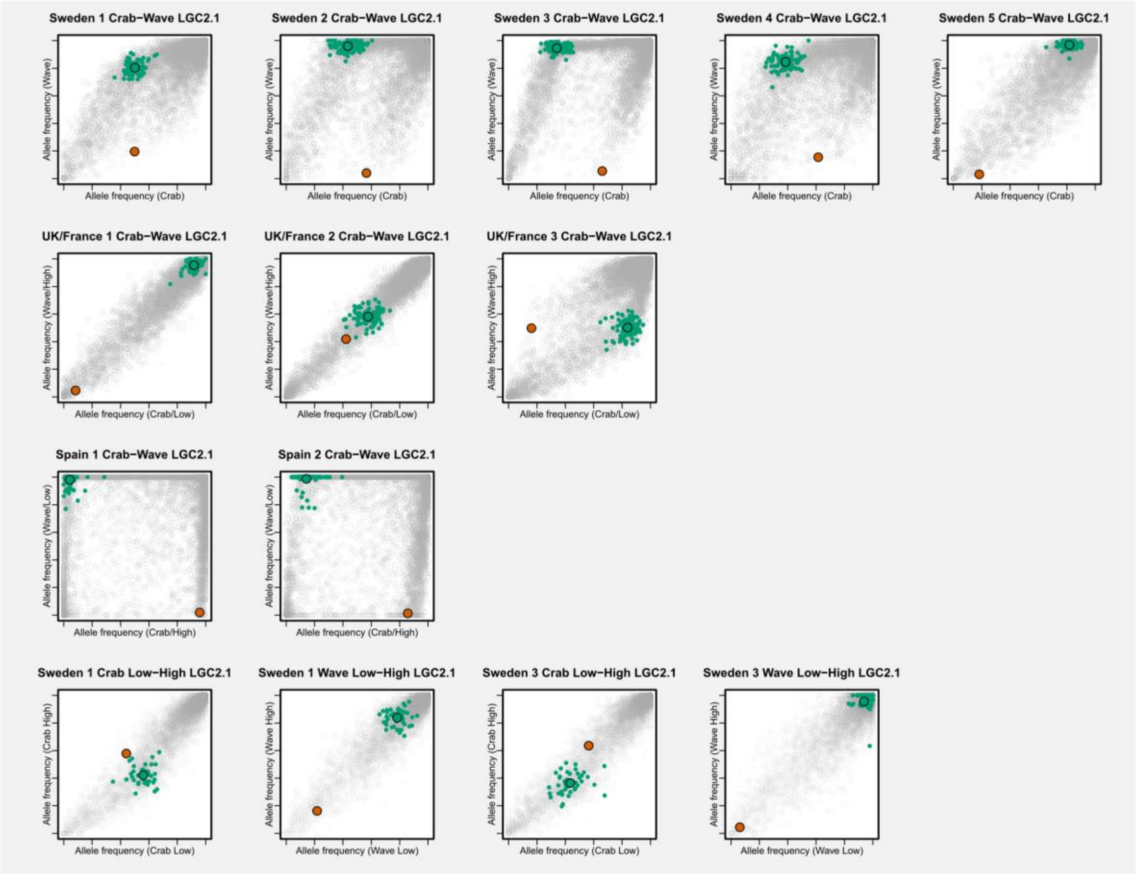

Sweden 1 Crab–Wave LGC2.1
Sweden 2 Crab–Wave LGC2.1
Sweden 3 Crab–Wave LGC2.1
Sweden 4 Crab–Wave LGC2.1
Sweden 5 Crab–Wave LGC2.1
Allele frequency (Wave)
Allele frequency (Crab)
UK/France 1 Crab–Wave LGC2.1
UK/France 2 Crab–Wave LGC2.1
UK/France 3 Crab–Wave LGC2.1
Allele frequency (Wave/High)
Allele frequency (Crab/Low)
Spain 1 Crab–Wave LGC2.1
Spain 2 Crab–Wave LGC2.1
Allele frequency (Wave/Low)
Allele frequency (Crab/High)
Sweden 1 Crab Low–High LGC2.1
Sweden 1 Wave Low–High LGC2.1
Sweden 3 Crab Low–High LGC2.1
Sweden 3 Wave Low–High LGC2.1
Allele frequency (Crab High)
Allele frequency (Crab Low)
Allele frequency (Wave High)
Allele frequency (Wave Low)

#### LGC6.1/2:

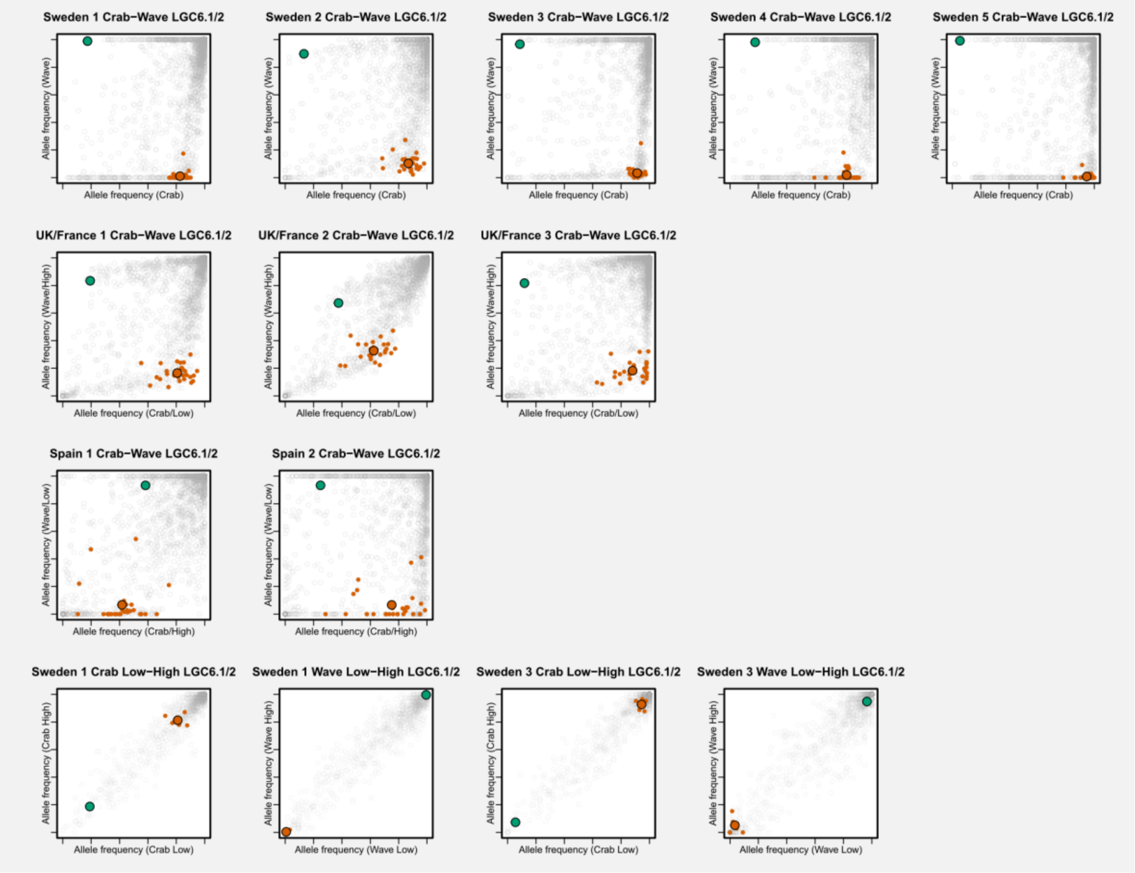


#### LGC7.1:

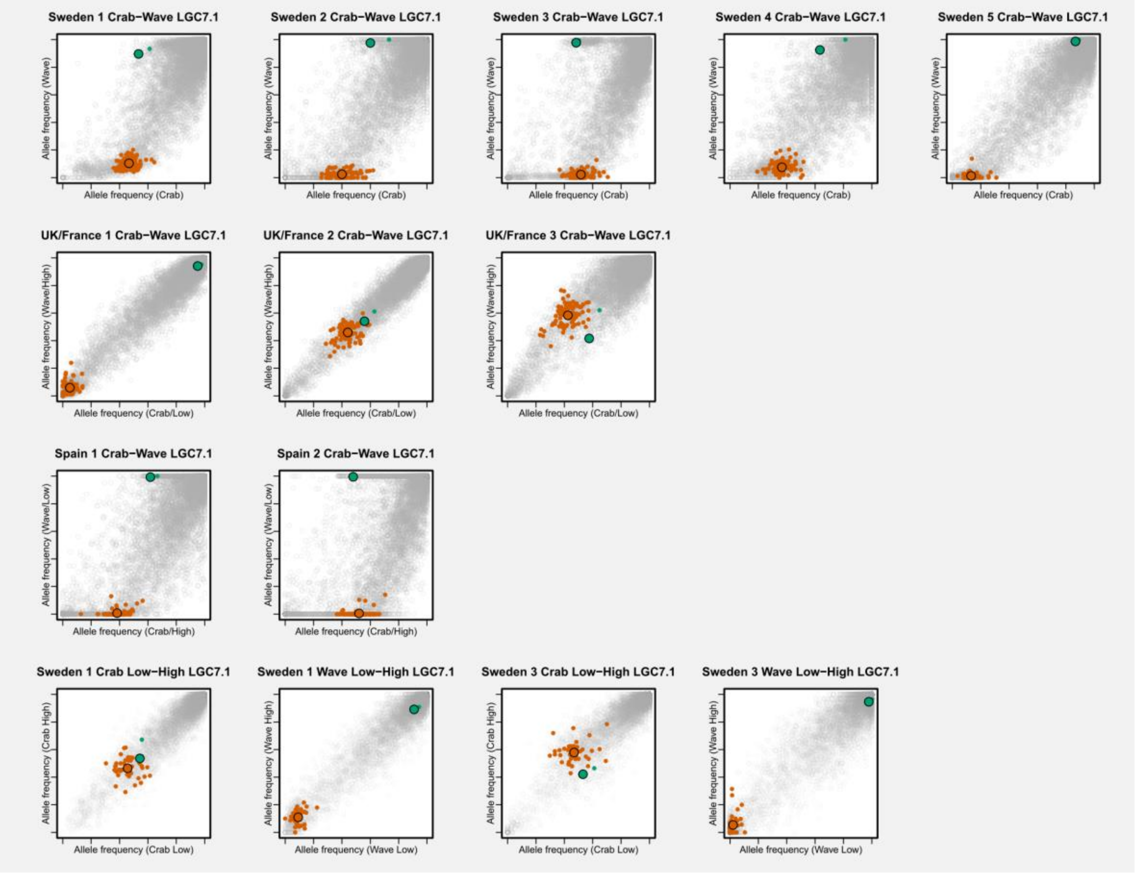

**LGC7.2:**

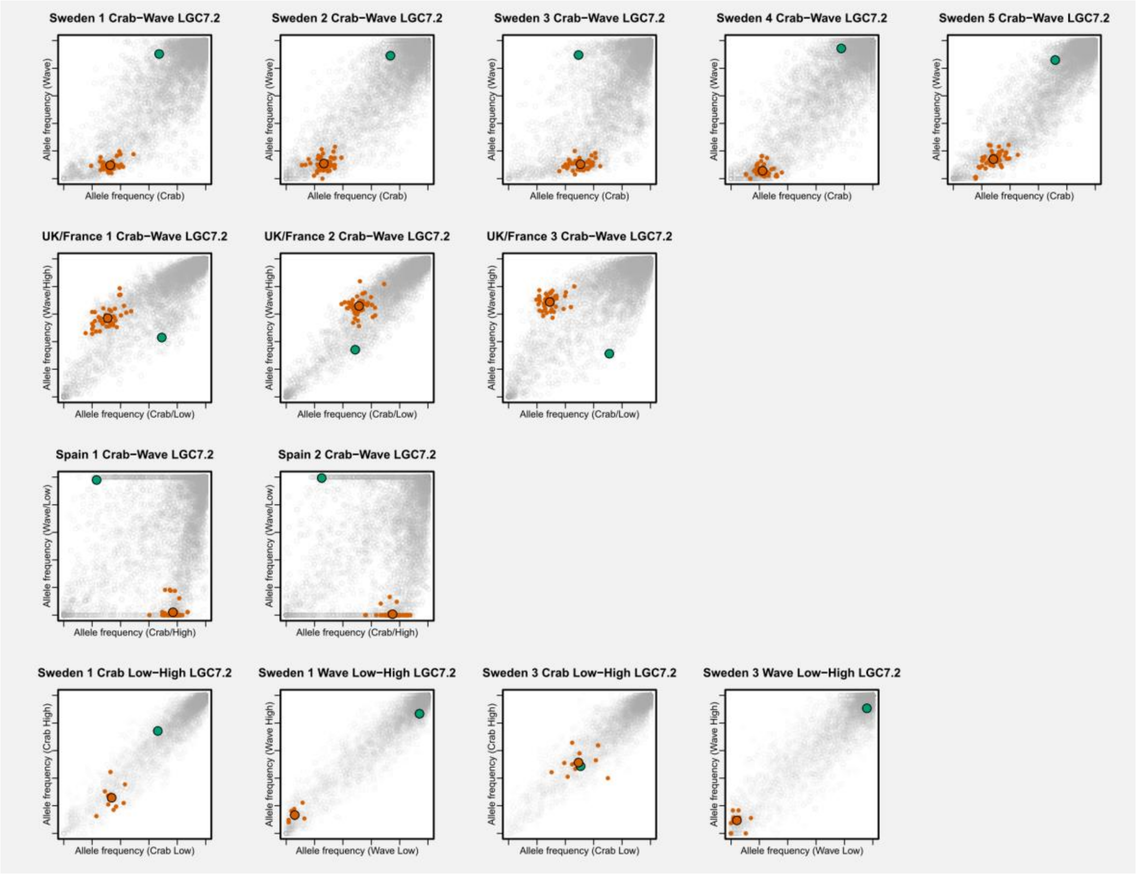

Sweden 1 Crab−Wave LGC7.2
Sweden 2 Crab−Wave LGC7.2
Sweden 3 Crab−Wave LGC7.2
Sweden 4 Crab−Wave LGC7.2
Sweden 5 Crab−Wave LGC7.2
Allele frequency (Wave)
Allele frequency (Crab)
UK/France 1 Crab−Wave LGC7.2
UK/France 2 Crab−Wave LGC7.2
UK/France 3 Crab−Wave LGC7.2
Allele frequency (Wave/High)
Allele frequency (Crab/Low)
Spain 1 Crab−Wave LGC7.2
Spain 2 Crab−Wave LGC7.2
Allele frequency (Wave/Low)
Allele frequency (Crab/High)
Sweden 1 Crab Low−High LGC7.2
Sweden 1 Wave Low−High LGC7.2
Sweden 3 Crab Low−High LGC7.2
Sweden 3 Wave Low−High LGC7.2
Allele frequency (Crab High)
Allele frequency (Crab Low)
Allele frequency (Wave High)
Allele frequency (Wave Low)


**LGC9.1:**

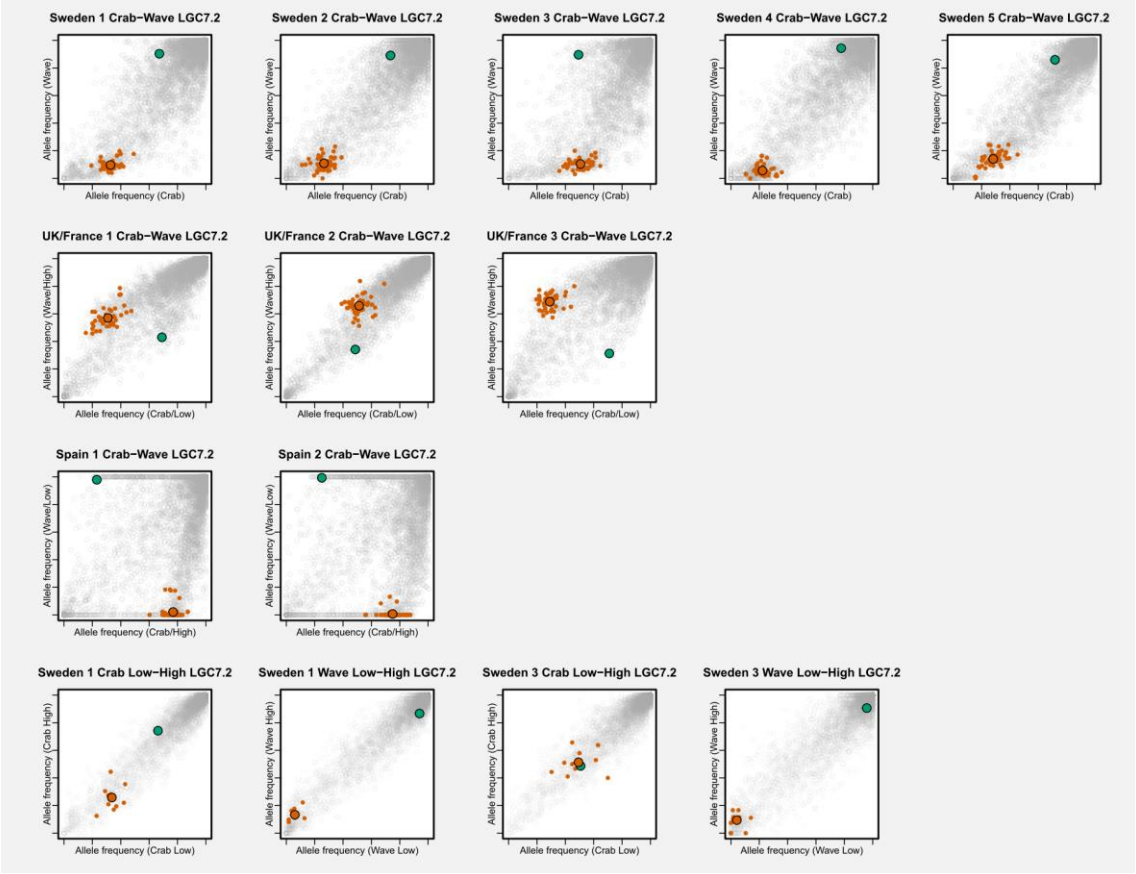

Sweden 1 Crab−Wave LGC7.2
Sweden 2 Crab−Wave LGC7.2
Sweden 3 Crab−Wave LGC7.2
Sweden 4 Crab−Wave LGC7.2
Sweden 5 Crab−Wave LGC7.2
Allele frequency (Wave)
Allele frequency (Crab)
UK/France 1 Crab−Wave LGC7.2
UK/France 2 Crab−Wave LGC7.2
UK/France 3 Crab−Wave LGC7.2
Allele frequency (Wave/High)
Allele frequency (Crab/Low)
Spain 1 Crab−Wave LGC7.2
Spain 2 Crab−Wave LGC7.2
Allele frequency (Wave/Low)
Allele frequency (Crab/High)
Sweden 1 Crab Low−High LGC7.2
Sweden 1 Wave Low−High LGC7.2
Sweden 3 Crab Low−High LGC7.2
Sweden 3 Wave Low−High LGC7.2
Allele frequency (Crab High)
Allele frequency (Crab Low)
Allele frequency (Wave High)
Allele frequency (Wave Low)

**LGC10.1:**

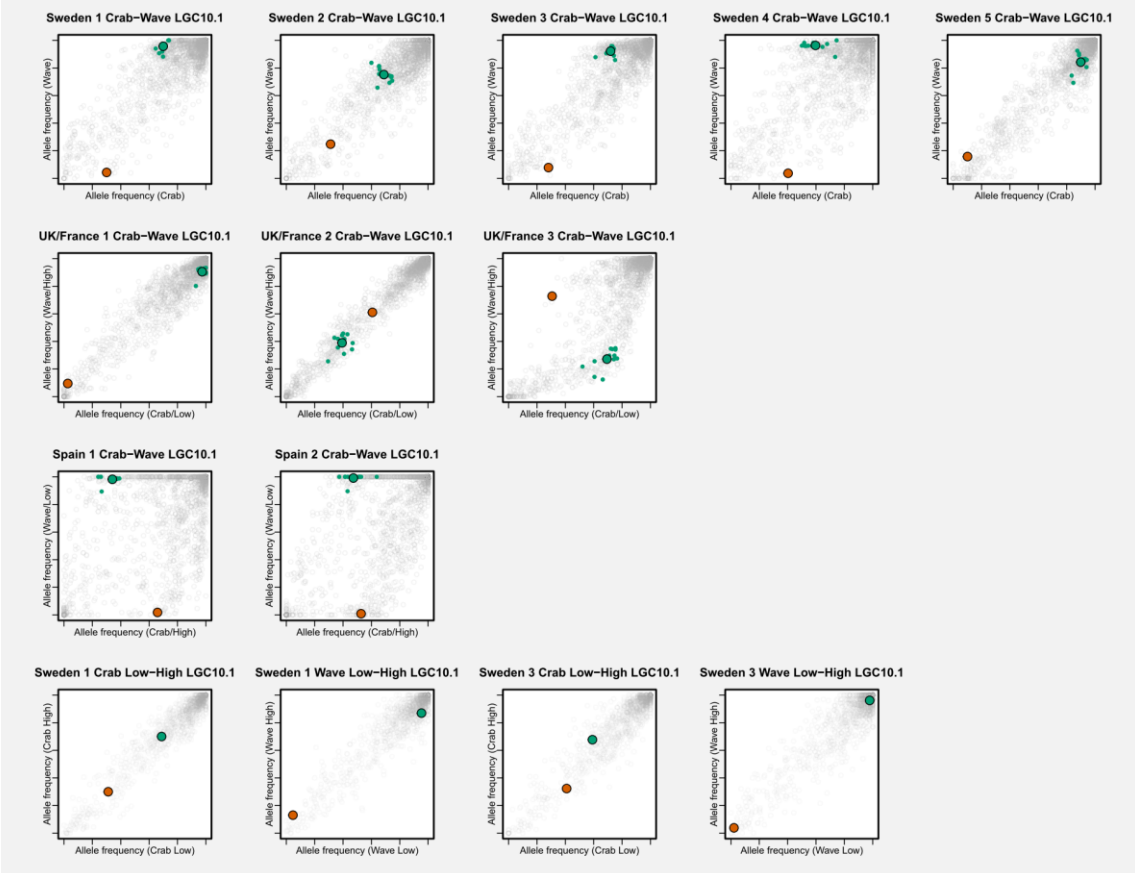


**LGC10.2:**

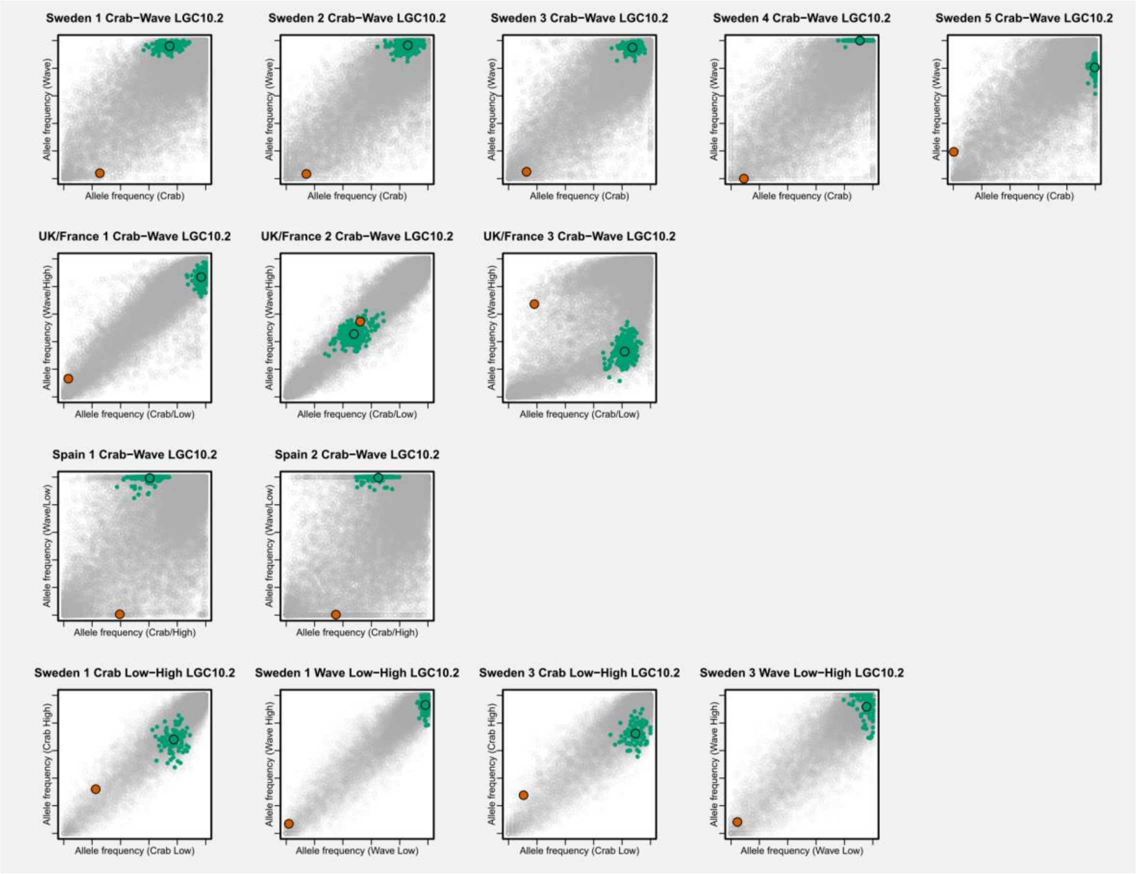

**LGC11.1:**

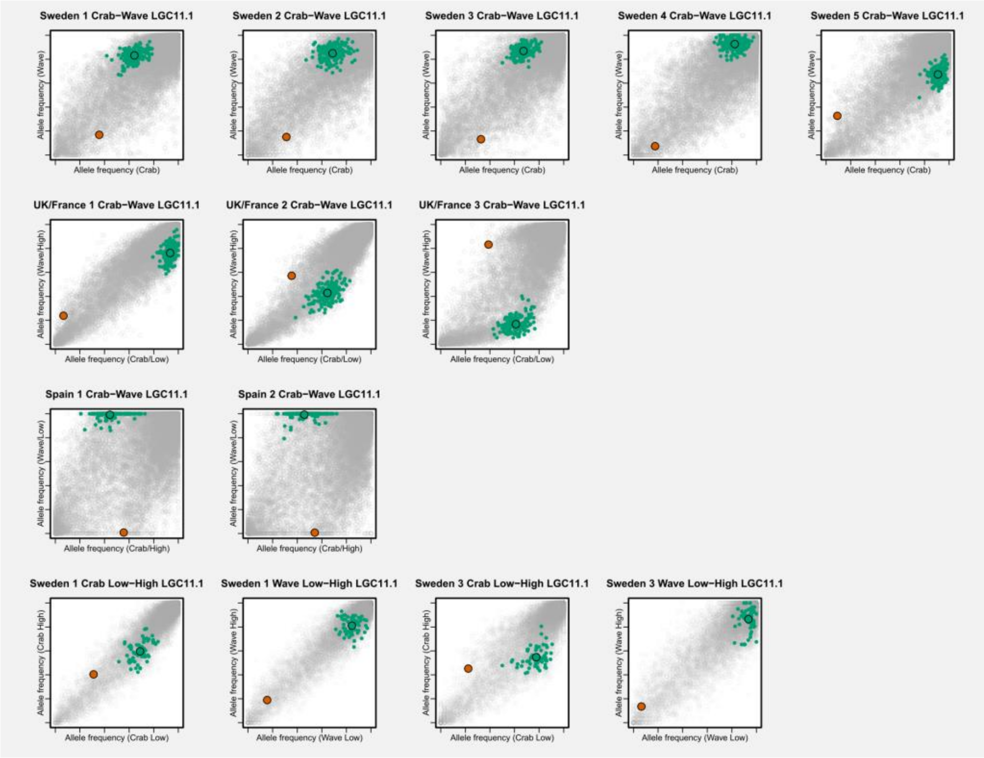


**LGC17.1:**

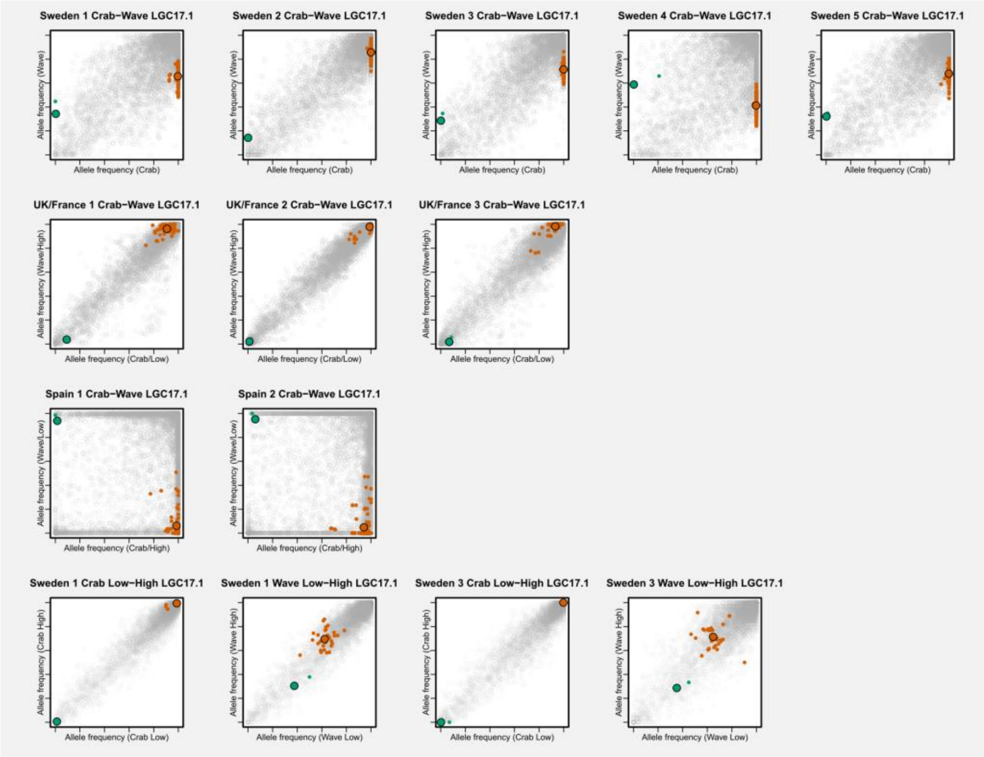


Fig. S3: Allele frequency plots for all inversion regions (also see Fig. 3 in the main text). For each inversion, we show 14 different pairs of samples. Axes range from 0 to 1. Each small point represents the frequency of the reference genome allele for a SNP in the inversion region. Coloured points represent arrangement-diagnostic SNPs (green: arrangement A; orange: arrangement B). Larger points with black borders indicate the frequencies of arrangement A and B obtained by averaging frequencies across arrangement-diagnostic SNPs.

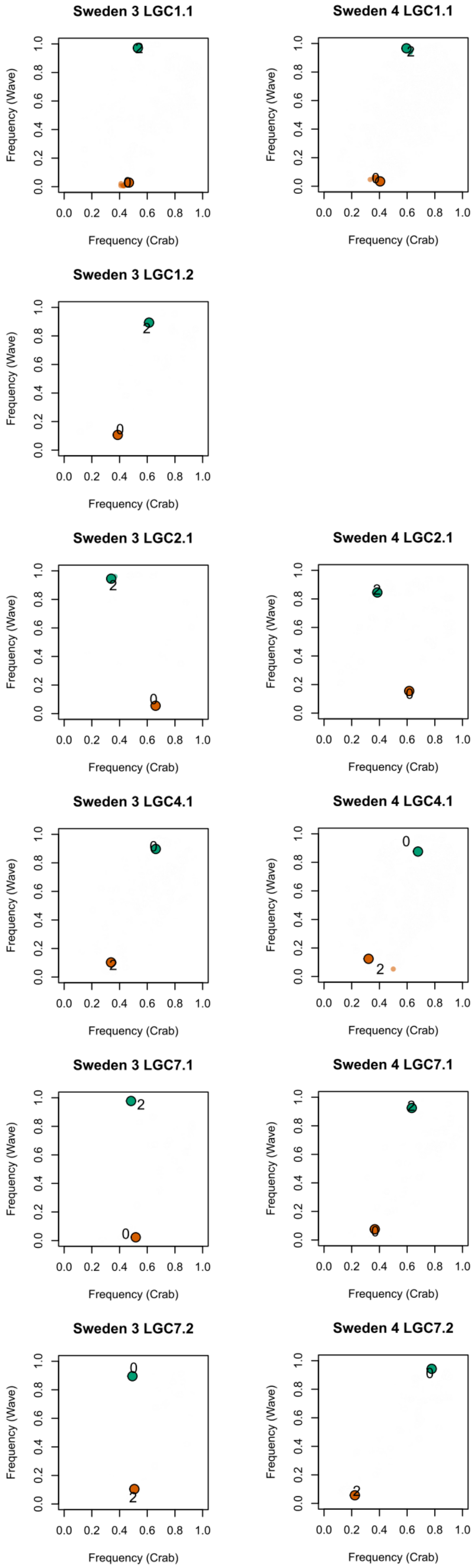
Sweden 3 LGC1.1
Sweden 4 LGC1.1
Sweden 3 LGC1.2
Sweden 3 LGC2.1
Sweden 4 LGC2.1
Sweden 3 LGC4.1
Sweden 4 LGC4.1
Sweden 3 LGC7.1
Sweden 4 LGC7.1
Sweden 3 LGC7.2
Sweden 4 LGC7.2
Frequency (Wave)
Frequency (Crab)
0.0 0.2 0.4 0.6 0.8 1.0

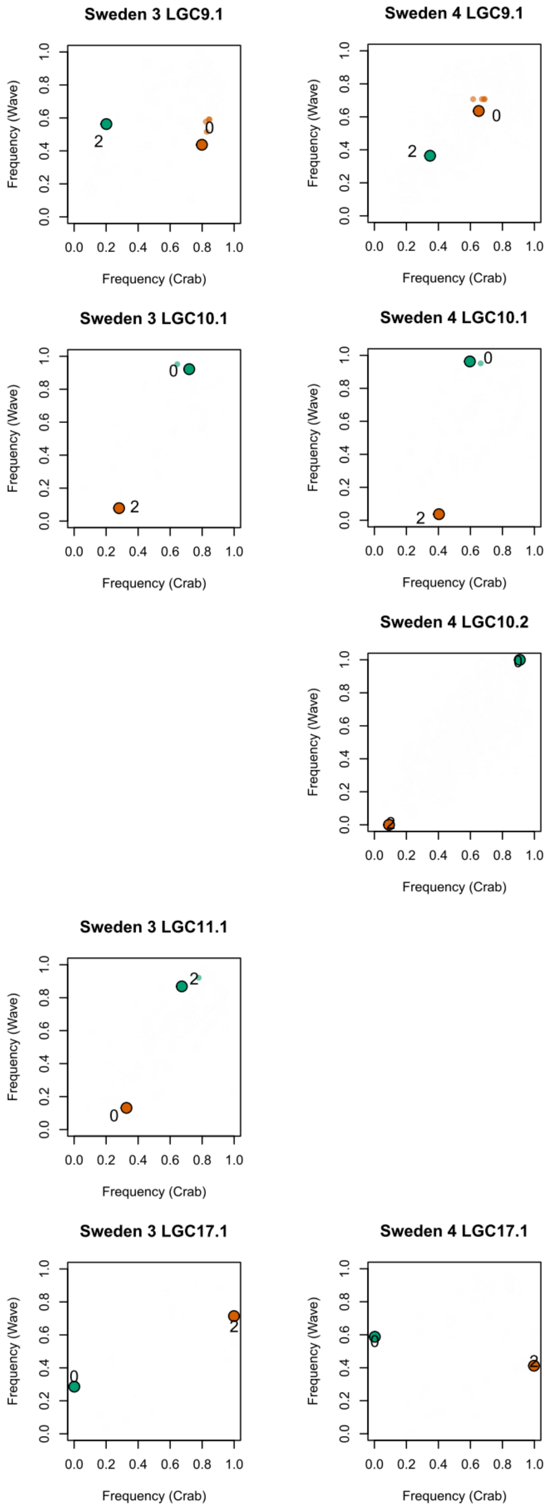


Fig. S4: Comparison between arrangement frequencies obtained from individual sequencing data from Westram et al. (2021) (indicated by the arrangement IDs used in that study, 0 and 2) and in the current study (dots; arrangement A: green, arrangement B: orange). Each row shows one inversion; the two columns are the two sampling locations that were included both in the current study and in Westram et al. (2021). Arrangement-diagnostic SNPs identified in the current study that were also included in the individual sequencing dataset are shown as smaller, semi-transparent dots (A-diagnostic SNPs: green; B-diagnostic SNPs: orange). For some inversions, arrangement frequency estimation was not possible for the individual sequencing data; in that case no plot is shown. The complex inversion (LGC6.1/2) is also not shown. This figure shows that the arrangements from the two studies can clearly be assigned to each other, that this assignment is consistent between the two locations included, and that frequency estimates are similar between studies.

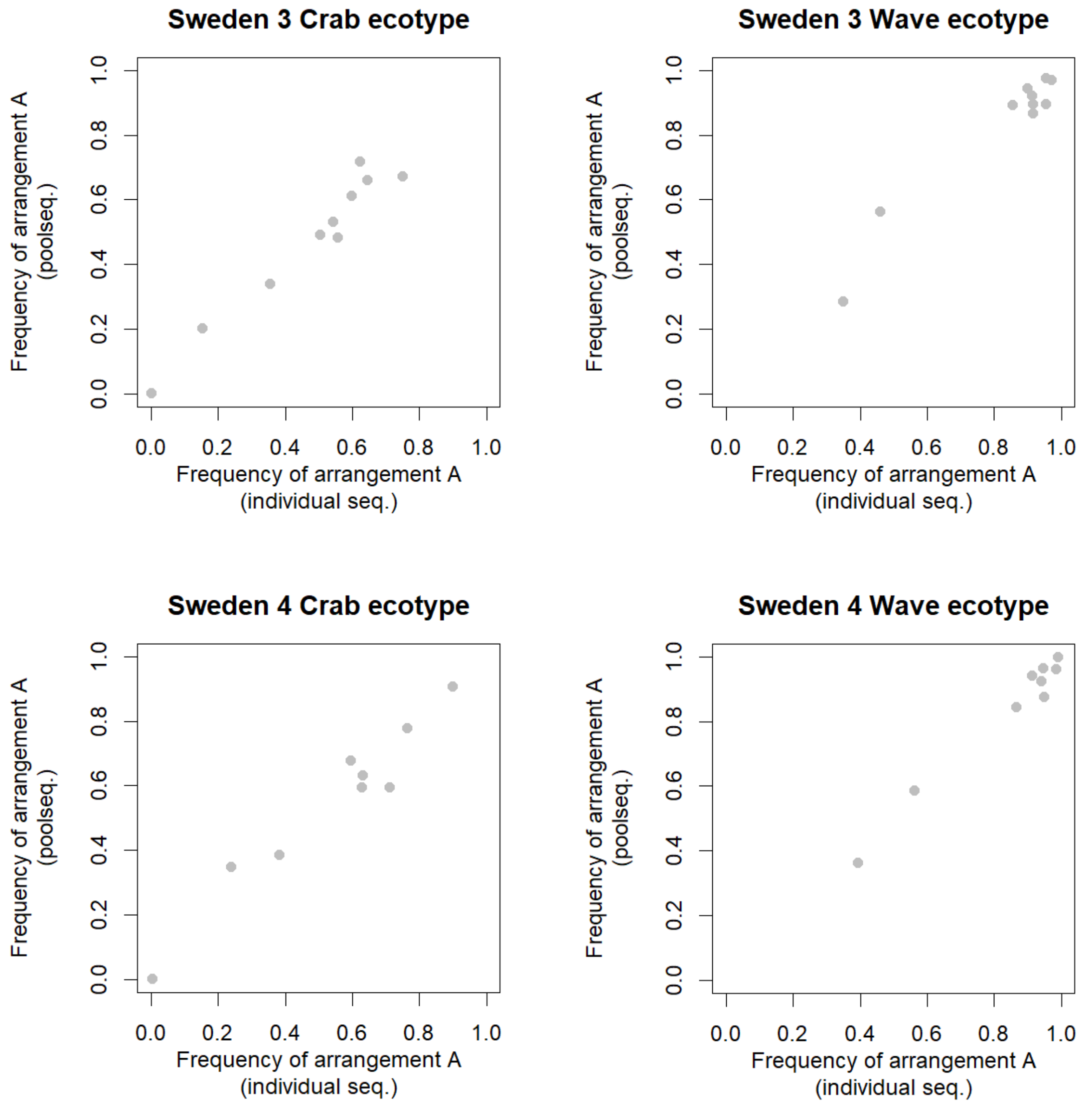


Fig. S5: Relationship between arrangement frequency estimates obtained with individual sequencing data (capture sequencing and linkage disequilibrium analysis) and pool-seq data (this paper) for the two ecotypes in two Swedish locations. Because different datasets and independent methods were used to identify arrangement frequencies in the two studies, we could not know a priori which of the two arrangements identified in the individual sequencing data (labelled arrangement 0 and 2) corresponded to arrangement A. Before making this plot we therefore assigned one of the two arrangements from the individual sequencing data to arrangement A, depending on which one was more similar in frequency; this was always obvious (see Fig. S4). We excluded the complex rearrangement LGC6.1/2 from this analysis. The correlation between frequencies from individual and pool-seq data was highly significant in all cases (Sweden 3 Crab ecotype: r=0.98; Sweden 3 Wave ecotype: r=0.97; Sweden 4 Crab ecotype: r=0.97; Sweden 4 Wave ecotype: r=0.99. All p<0.0001).

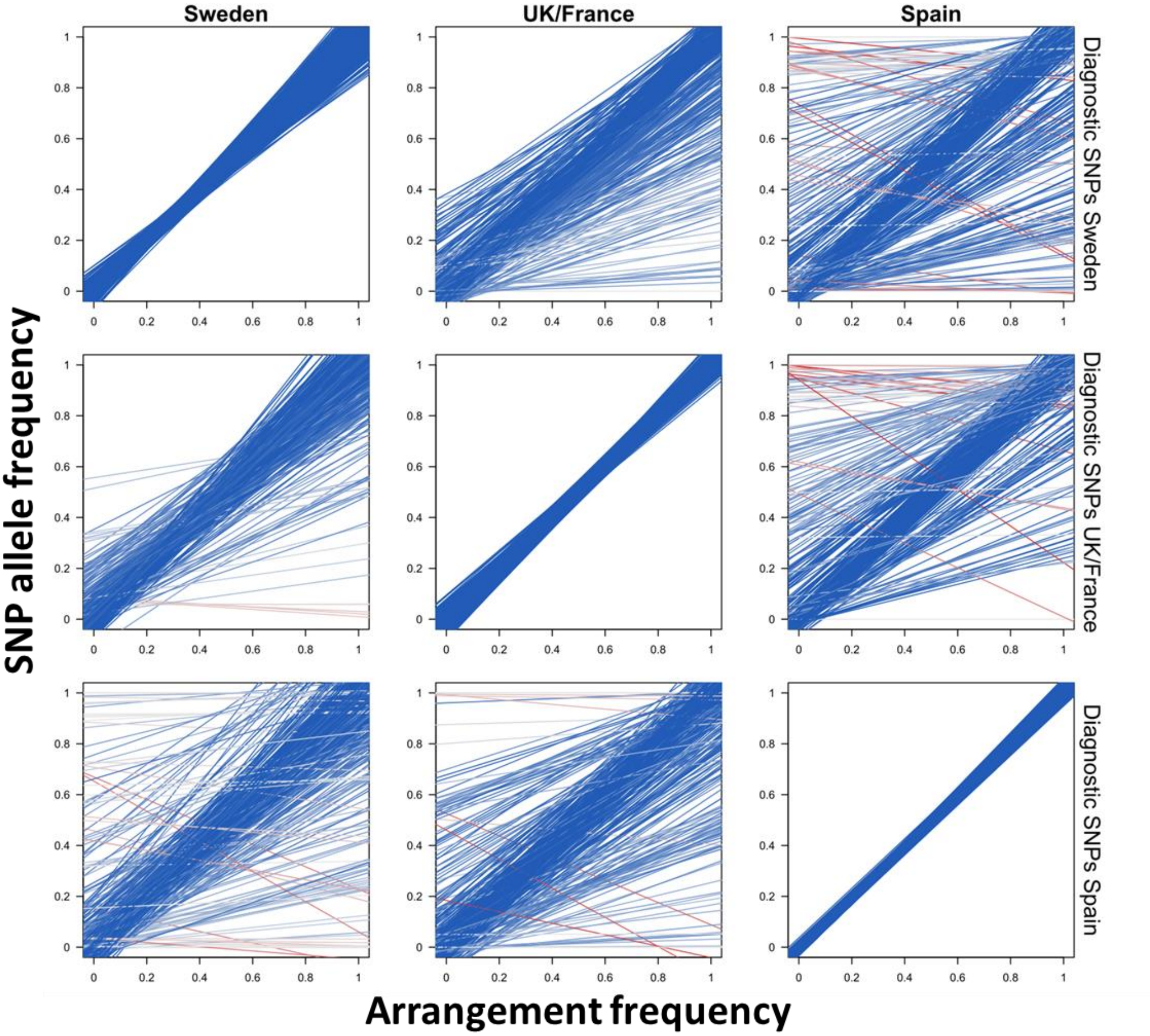


Fig. S6: Patterns of locally arrangement-diagnostic SNPs in other geographical areas for the LGC9.1 inversion. Each line represents the regression of SNP allele frequency against the frequency of arrangement A for one SNP. The geographical area where the SNPs are arrangement -diagnostic is indicated on the right (SNPs can be arrangement-diagnostic in multiple countries, so they can occur in multiple rows); regression lines always show the allele that is positively associated with arrangement A in this geographical area. Line colour indicates Pearson’s correlation coefficient (r), from negative (red) via 0 (grey) to positive (blue).

### *Supplementary Tables*

Table S1: Samples included in this study. The samples were the same as in Morales et al. (2019), but we use different labelling.

| Location ID this study | Location ID in Morales et al. 2019 | Country | Locality | Latitude | Longitude | Samples |
|---|---|---|---|---|---|---|
| Sweden 1 | SWn5 | Sweden | Arsklovet | 58.832798 | 11.134298 | Crab High, Crab Low, Wave High, Wave Low |
| Sweden 2 | SWn1 | Sweden | Jutholmen | 58.87112 | 10.985244 | Crab, Wave |
| Sweden 3 | SWn2 | Sweden | Ramso | 58.831282 | 11.0669 | Crab High, Crab Low, Wave High, Wave Low |
| Sweden 4 | SWn4 | Sweden | Salto | 58.866114 | 11.134091 | Crab, Wave |
| Sweden 5 | SWs | Sweden | Öckerö | 57.71175 | 11.631305 | Crab, Wave |
| UK/France 1 | Uke | UK | Thornwick | 54.1328 | -0.1129 | Crab Low, Wave High |
| UK/France 2 | Ukw | UK | Anglesey | 53.2999 | -4.6795 | Crab Low, Wave High |
| UK/France 3 | Fr | France | Pointe de Primel | 48.718462 | -3.820313 | Crab Low, Wave High |
| Spain 1 | SPn | Spain | Burela | 43.6763 | -7.3679 | Crab High, Wave Low |
| Spain 2 | SPs | Spain | Silleiro | 42.111684 | -8.898521 | Crab High, Wave Low |

### Supplementary Text

#### Illustrative simulation of allele frequency patterns in inversion regions

For Fig. 1 in the main text, we performed simple simulations illustrating how parallelograms emerge in allele frequency plots. We did not simulate population dynamics but instead only aimed at generating semi-realistic allele frequencies in populations connected by gene flow containing inversion polymorphism.

The simulations for Fig. 1C) and D) contained 400 SNPs variable in arrangement A and 400 SNPs variable in arrangement B. Allele frequencies for each SNP were randomly sampled from a beta distribution with shape parameters 0.1, 0.1 for the arrangement where it was variable, and for the other arrangement randomly set to 0 or 1. The shape of the beta distribution ensures that many SNPs are essentially fixed different between arrangements.

We assumed that the allele frequency within an arrangement was the same in population 1 and 2. We varied two parameters, the frequency of arrangement A in population 1 ($p_{A,1}$) and the frequency of arrangement A in population 2 ($p_{A,2}$). We calculated the allele frequency of each SNP in each population (using $p_{A,1}$, $p_{A,2}$ and the frequency of each SNP within each arrangement). Finally, we introduced sampling noise by randomly sampling 100 copies of each SNP in each population and then re-calculating allele frequencies. We plotted the resulting SNP allele frequencies in population 2 against those in population 1. We considered SNPs with an underlying frequency of less than 0.01 in one arrangement and higher than 0.99 in the other arrangement as "arrangement-diagnostic" (highlighted in Fig. 1 C) and D)).

The simulations for the collinear region shown in Fig. 1A) were done similarly as for the regions with inversion polymorphism, except that we assumed a single arrangement with 800 SNPs.

#### Determination of arrangement-diagnostic SNPs

We performed the following steps for each inversion region (We illustrate each step for inversion LGC9.1 and the location Sweden 3):

**Step 1. Visual inspection of parallelograms**

We plotted population 2 (here: Wave population) allele frequencies against population 1 (here: Crab population) allele frequencies for each location separately to generate plots analogous to Fig. 1C/D in the main text. We checked the allele frequency plots for the presence of parallelograms. All steps for the determination of arrangement-diagnostic SNPs were automated (see below); however, our approach relies on at least subtle arrangement frequency differences between populations in multiple locations, so that it makes sense to first visually check whether this requirement is fulfilled. In our dataset, each inversion region showed parallelograms in multiple locations, so we proceeded with the following steps for all inversions.

If there was a parallelogram only in a single location, one could identify arrangement-diagnostic SNPs based on that location only (by determining the SNPs in the parallelogram corners based on some threshold) and then check their frequencies in other locations to estimate arrangement frequencies where there is no strong frequency difference between local populations. This is likely to work reliably on a small geographical scale, but might fail on larger scales as the diagnostic SNPs in this case have not been ensured to be diagnostic across many locations.

If there was no parallelogram in any location, it could still be possible that inversion polymorphism exists but frequency differences are too subtle or the data are too noisy to generate visible patterns. If the presence of an inversion is suspected based on other results, it could therefore still make sense to attempt the approach below, but to carefully examine all intermediate steps to check whether they are consistent with the presence of an inversion.

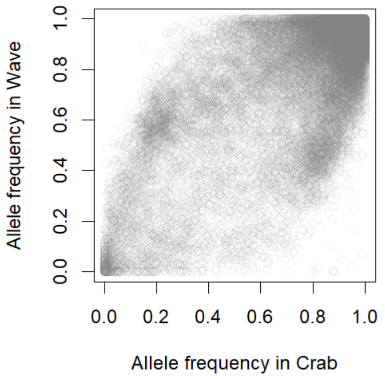


Fig. T1: Allele frequency plot including all SNPs in the inversion region for the location Sweden 3. This plot shows a clear parallelogram, with the SNPs (nearly) fixed different between arrangements forming the corners with a higher density of points.

#### 2. Finding candidate arrangement-diagnostic SNPs

The differentiation ($F_{ST}$) between the local populations (e.g. Crab and Wave ecotype) in an inversion region is typically highest for the arrangement-diagnostic SNPs. To identify candidate arrangement-diagnostic SNPs we therefore selected those SNPs that fell above the 80% $F_{ST}$ quantile in at least 50% of the geographical locations.

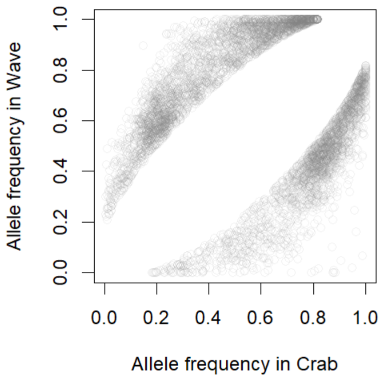


Fig. T2.1: High-$F_{ST}$ SNPs determined for the location Sweden 3.

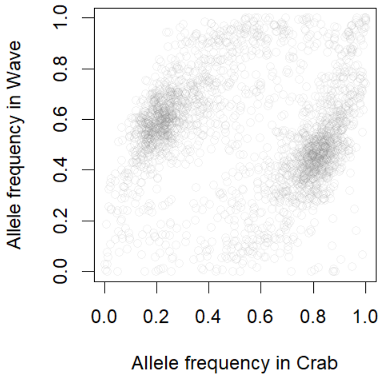


Fig. T2.2: Candidate inversion-diagnostic SNPs obtained by retaining only those SNPs that showed a high Crab-Wave $F_{ST}$ in at least 50% of the locations, again shown for Sweden 3. It can be seen that this step enriches for SNPs in the parallelogram corners, but retains some SNPs that do not seem to be inversion-diagnostic in the focal location.

**Step 3. Confirmation of candidate arrangement-diagnostic SNPs using covariation between ecotypes and locations**

Arrangement-diagnostic SNPs are expected to not only show high $F_{ST}$ between local populations, but also to strongly co-vary with each other across different samples due to consistent LD with the inversion. We therefore ran a PCA on all samples included in the analysis, using only the frequencies of the candidate arrangement-diagnostic SNPs identified in step 2. The first principal component then arranges samples roughly by arrangement frequency, and the SNPs most strongly contributing to this axis are most strongly associated with the arrangements. SNPs where the reference allele is associated with one arrangement will make a positive contribution to PC1; SNPs where the reference allele associated with the other arrangement will make a negative contribution. We arbitrarily labelled the arrangement associated with positive values on PC1 "arrangement A" and the one associated with negative values "arrangement B". To include only the SNPs most strongly associated with the inversion, we calculated the 90% quantile of the absolute values of the PC1 loadings, and only included those SNPs in our set of diagnostic SNPs that fell above this threshold. Those with positive loadings were stored as A-diagnostic SNPs and those with negative loadings as B-diagnostic SNPs.

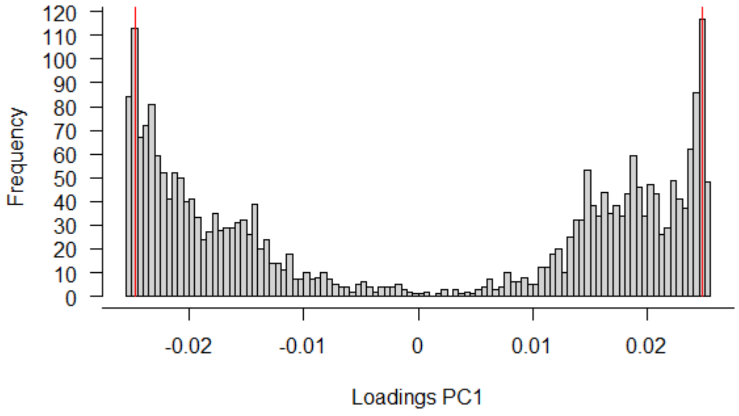


Fig. T3.1: PC1 loadings from a PCA across all samples, using only candidate inversion-diagnostic SNPs. The SNPs most strongly associated with each other, and with the arrangement, have extreme values. We therefore considered those SNPs inversion-diagnostic that were located about the 90% quantile of the distribution of absolute loading values (i.e. to the left of the left red line [B-diagnostic SNPs] or to the right of the right red line [A-diagnostic SNPs]).

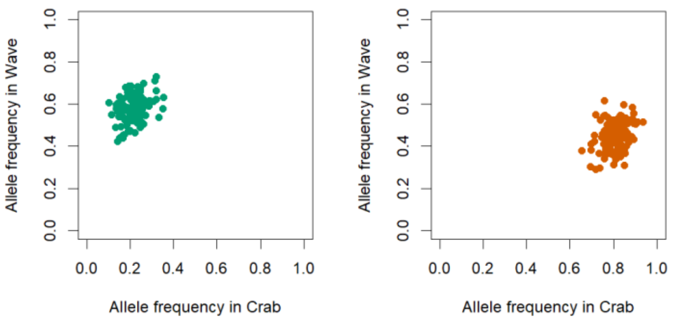


Fig. T3.2: Allele frequencies of final arrangement-diagnostic SNPs in Sweden 3 (left: A-diagnostic; right: B-diagnostic). A comparison with the plots from earlier steps demonstrates that these SNPs are clearly concentrated in the parallelogram corners.

### *References*

Westram, A. M., R. Faria, K. Johannesson, and R. Butlin. 2021. Using replicate hybrid zones to understand the genomic basis of adaptive divergence. Molecular Ecology 30:3797–3814.
